# Single-nucleus and spatial transcriptomic analysis identified molecular features of neuronal heterogeneity and distinct glial responses in Parkinson’s disease

**DOI:** 10.1101/2024.07.28.605055

**Authors:** Sooyeon Yoo, Kwanghoon Lee, Junseo Seo, Hwisoo Choi, Seong-Ik Kim, Junyoung Chang, Yu-Mi Shim, Junil Kim, Jae-Kyung Won, Sung-Hye Park

## Abstract

The heterogeneity of Parkinson’s disease (PD) is increasingly recognized as an important aspect of understanding the disorder. Among the factors contributing to this heterogeneity, ethnic differences are primary sources, significantly influencing the likelihood of PD developing and its initial symptoms’ nature. While there have been numerous reports related to PD in East Asia, there has been a lack of contribution from single-cell (or nucleus) transcriptome studies, which have been making significant contributions to understanding PD. In this study, a total of 33,293 nuclei obtained from the substantia nigra (SN) of confirmed pathological PD and control patients in South Korea were profiled, revealing 8 different cell types through cluster analysis. Monocle-based pseudotime analysis identified two disease-associated trajectories for each astrocyte and microglia and identified genes that differentiate them. Interestingly, we uncovered the inflammatory intervention in the early PD-associated transition in microglia and identified the molecular features of this intermediate state of microglia. In addition, gene regulatory networks (GRNs) based on TENET analysis revealed the detrimental effect of an *HSPA5-*led module in microglia and *MSRB3- and HDAC8-* led modules specifying the two different astrocyte trajectories. In SN neurons, we observed population changes, a decrease in dopaminergic and glutamatergic neurons and a proportional increase in GABAergic neurons. By deconvolution in spatial transcriptome obtained the PD sample, we confirmed spatiotemporal heterogeneity of neuronal subpopulations and PD-associated progressive gliosis specific to dopaminergic nuclei, SN and ventral tegmental areas (VTAs). In conclusion, our approach has enabled us to identify the genetic and spatial characterization of neurons and to demonstrate different glial fates in PD. These findings advance our molecular understanding of cell type-specific changes in the progression of Korean PD, providing an important foundation for predicting and validating interventions or drug effects for future treatments.

## Introduction

Parkinson’s disease (PD) is the second most common neurodegenerative disorder, characterized by the abnormal accumulation of alpha-synuclein (α-Syn) leading to the formation of Lewy bodies and the loss of dopaminergic neurons in the substantia nigra (SN) of the midbrain. As with other neurodegenerative diseases, early detection of PD is challenging, as symptoms manifest only after significant dopaminergic cell loss has occurred. Clinical diagnosis is based on observable movement problems like bradykinesia, rigidity, tremor, and postural instability. However, the pathological features of PD mentioned above can only be confirmed through a comprehensive pathological examination of postmortem brain tissue.

Genetic studies, including pedigree analysis, whole-exome sequencing, and genome-wide association studies (GWAS) have provided valuable insights into the genetic factors associated with PD, some of which are considered causal genes and potential therapeutic targets [1,2]. However, these genes exhibit diverse genetic variants across different races and the same variant has different impacts. For example, the G2019S variant of LRRK2, known for increasing kinase activity, is rare in some racial groups [3–6], while G2385R and R1628P are commonly found in Asian populations and associated with increased risk of PD [7,8]. Various LRRK2 kinase inhibitors are currently in clinical trials [9,10], and it is now imperative to consider interethnic heterogeneities, which are believed to explain clinical disparities [11–13] and to move towards personalized treatments.

To gain a better understanding of sporadic PD, recent GWAS studies have expanded to include Asian populations in addition to those previously limited to Europe and the United States [14–18]. By comparing and analyzing the results from Western and Eastern populations, more clearly the interethnic differences in PD were observed. A notable example is the recent GWAS study on the Korean population, which highlighted differences in genetic contributions to sporadic PD based on race and gender [19,20]. Efforts are also being made on a worldwide level to include diverse populations in genetic research initiatives, although it is still limited and requires more cooperation. Beyond just genetic variations, it is crucial to conduct diverse modalities of genetic research to understand how racial differences may impact the conditions experienced by PD patients.

Single-cell transcriptome sequencing has been actively applied to PD research in recent years, as it has the great advantage of observing gene expression changes in a wide variety of cell types in the human brain [21–27]. These studies have utilized samples of nuclei extracted from the midbrain or SN region of PD patients or samples enriched for dopaminergic neurons, which are the primary neuronal populations affected in PD. However, determining consistent genetic features across various PD data sources and integrating the multifaceted information to resolve outstanding questions about the disease remains challenging. In particular, these studies have been conducted primarily in white participants or have not distinguished between racial groups, and there is a lack of results from studies conducted in the Asian population.

Considering the heterogeneity of PD itself, it is important to take into account the pathological subclassification of PD and to use homogeneous groups for analysis to reduce variation and improve the accuracy of the research. In our pathological diagnoses, we recognize that many patients with no neurodegeneration and no significant cognitive impairment turned out to have primary age-related tauopathy (PART), which is consistent with previous reports [28,29]. Therefore, for the selection of the control group in this study, we specifically chose to use those cases confirmed with PART pathology. By using such a consistent pathological control group, we can expect the following advantages. First, the pathological process precedes the clinical stage of PD, so the pathological confirmation evaluates the undesirable progression of the disease in control cases. Second, patients with neurodegenerative diseases experience aging along with pathology, so it is important to distinguish natural aging from disease progression. Therefore, using individuals who do not progress to disease despite normal aging as a control group allows for a more accurate distinction between pathological changes and normal aging.

In this study, we identified eight different cell types, including neurons, oligodendrocytes, oligodendrocyte precursor cells (OPCs), astrocytes, microglia, endothelial cells, and T cells in the SN, and had a remarkable number of differential expression genes (DEGs) in PD compared to controls. We performed independent clustering within each cell type and confirmed high heterogeneity in SN neurons and found population changes. The number of cells in dopaminergic neuronal clusters decreased significantly in PD, and the glutamatergic clusters also showed a decreasing trend, but interestingly GABAergic populations were relatively less susceptible. In addition, the spatial transcriptome results revealed distinct spatial differences in neuronal subtypes majorly between the SN and VTA, which were strongly affected by PD, and the dorsolateral VTA, which was less affected. In the astrocyte and microglial subsets, the change from control to PD was more obvious than the difference between subclusters. Therefore, we performed a Monocle trajectory analysis to observe the expression changes across the continuous pseudotime relevant to the PD progression. Two trajectories related to PD were identified for each astrocyte and microglia, with the main differences attributed to inflammation, different ER stress responses, or abnormal reactivities. Notably, early involvement of inflammatory response was observed in microglial cells. Thus, by identifying various cell subtypes appeared during the progression of PD and confirming their genetic and spatial features, we were able to better explain the pathophysiological landscape changes. Furthermore, these results are expected to provide an important basis for future studies identifying pathophysiological function dependent on cell types and discovering important pathological markers that determine accurate PD diagnosis and disease severity, which may eventually become therapeutic targets.

## Methods

### Sample selection

Samples were obtained from brain donations to the Seoul National University Hospital Brain Bank (SNUH-BB). Following approval from the Institutional Review Board (IRB, approval number 1808-087-966), autopsies focusing only on the brain were conducted at SNUH-BB according to standardized protocols. Formalin-fixed, paraffin-embedded (FFPE) blocks were prepared and used for histopathologic analysis, employing techniques such as hematoxylin and eosin (H&E), Luxol fast blue (LFB), immunohistochemistry (IHC) [30]. For snRNA-seq analysis, four cases of PART and four cases of definite PD cases were selected. These cases were confirmed by three different neuropathologists (S-I K, J-K W, and S-H P) based on diagnostic criteria [31,32]. The median age was 74 years, with an equal male-to-female ratio, and a median postmortem interval of 9.25 hours. The PD cases had a median Braak Lewy body stage of 6 while PART cases had a stage of 0. Detailed sample information is provided in Supplementary Table 1. We carefully verified that there were no other pathological features besides PD in the midbrain. For spatial transcriptomics using 10X Visium V2, a case of PD at LB Braak stage 6 was selected. This individual developed PD at the young age of 45, allowing us to observe the pure influence of the disease without interference from other neurodegeneration conditions.

### Single nuclei isolation and sequencing library preparation

For snRNA-seq, we collected 30-50 mg of substantia nigra tissue from fresh frozen midbrain transverse sections, ensuring that the red nucleus was not included and that the entire region affected by PD was captured by an experienced pathologist (S-I K). The nuclear isolation protocol was implemented with some modifications from the 10x Genomics nuclei preparation protocol (#CG000124). The extracted nuclei were stained with 7-AAD (Invitrogen #00-6993-50) and the positive nuclei were isolated by fluorescence-activated cell sorting (FACS). Library preparation was performed according to the manufacturer’s instructions for the Chromium Next GEM Single Cell 3’ Reagent Kits v3.1 (Dual Index, #CG000315). Sequencing was performed on the Illumina Hi-seqX platform. Sequencing data from the FASTQ file was aligned using the default pipeline of 10x Genomics Cell Ranger 5.0.1. The reference genome for alignment and annotation was the human genome (GRCh38). After this process, a total of 33,293 cells were retained.

### snRNA data analysis pipeline

Data from eight samples were processed using the Seurat package (v4.0.3) in R [33]. Cells with a mitochondrial ratio less than 0.2 or that have UMI counts under 200 deemed to be of low quality, were excluded. Genes related to sex and mitochondria, which were considered to negatively impact the analysis, were removed. Genes that have total UMI counts under 3 were also removed. Genes related to ribosome, hemoglobin, platelet, and cell cycle were detected in the data, but due to the potential relevance to neurodegenerative diseases, they were not intentionally removed from the analysis due to a lack of evidence [34–36]. Following the quality control, count matrices were normalized. 2000 highly variable genes (HVGs) were used for principal component analysis (PCA). We removed doublets using the DoubletFinder package in R, applied individually to each sample [37]. The selection criterion for the pK value was the maximization of the bimodality coefficient. To cluster the nuclei by cell types, the data from different samples were integrated based on anchor cells implemented in Seurat [38]. Graph-based clustering with resolution 1 was then performed on a reduced principal dimension, resulting in 27 clusters. We deleted cluster 22, as it shows both oligodendrocytes and microglia marker genes. As a result, a total of 1533 cells were removed, 31760 cells remained.

### Cell type detection and DEG analysis

To identify cell types for each cluster, we utilized marker genes by comparing each cluster with all the other cells using Wilcoxon’s rank sum test with a log fold change threshold of 0.2, minimum expressed cell percentage of 0.01, and false discovery rate (FDR) threshold of 0.05. We categorized 26 clusters into 8 cell types based on the enriched marker genes. Next, we identified differentially expressed genes (DEGs) between CON and PD within each cell type using the same parameters as for the cell type marker gene identification. We excluded T cells and combined mural and endothelial cells for the DEG analysis.

We further clustered each cell type using the same procedure including 2,000 HVGs detection, PCA, and graph-based clustering. For accurate neuronal sub-cell type annotation, we integrated our neuron cells with public data containing dopaminergic neurons using the LIGER method implemented in Seurat Wrapper R library with default parameters [25,39].

### Evaluating the Overlap Between GWAS Genes and DEGs

From literature [14,16,20,40–45] and Gene4PD database [46], we collected 550 PD GWAS genes. For the Gene4PD database, we used an adjusted *p*-value threshold of 0.05. Hypergeometric test performed for intersection of PD GWAS genes and 12 DEG groups (upregulated and downregulated in PD across 6 cell types, as shown in Fig. 1G). Pseudogenes, long noncoding RNA genes, and microRNA genes, were excluded, resulting in a total pool of 29,635 genes for the hypergeometric test. The hypergeometric test was also performed in the same way with PD GWAS genes and spatial DEGs with log-fold change greater than 1 and adjusted *p*-value threshold of 0.05. (Supplementary Table 2_3 and 3)

**Figure 1.**
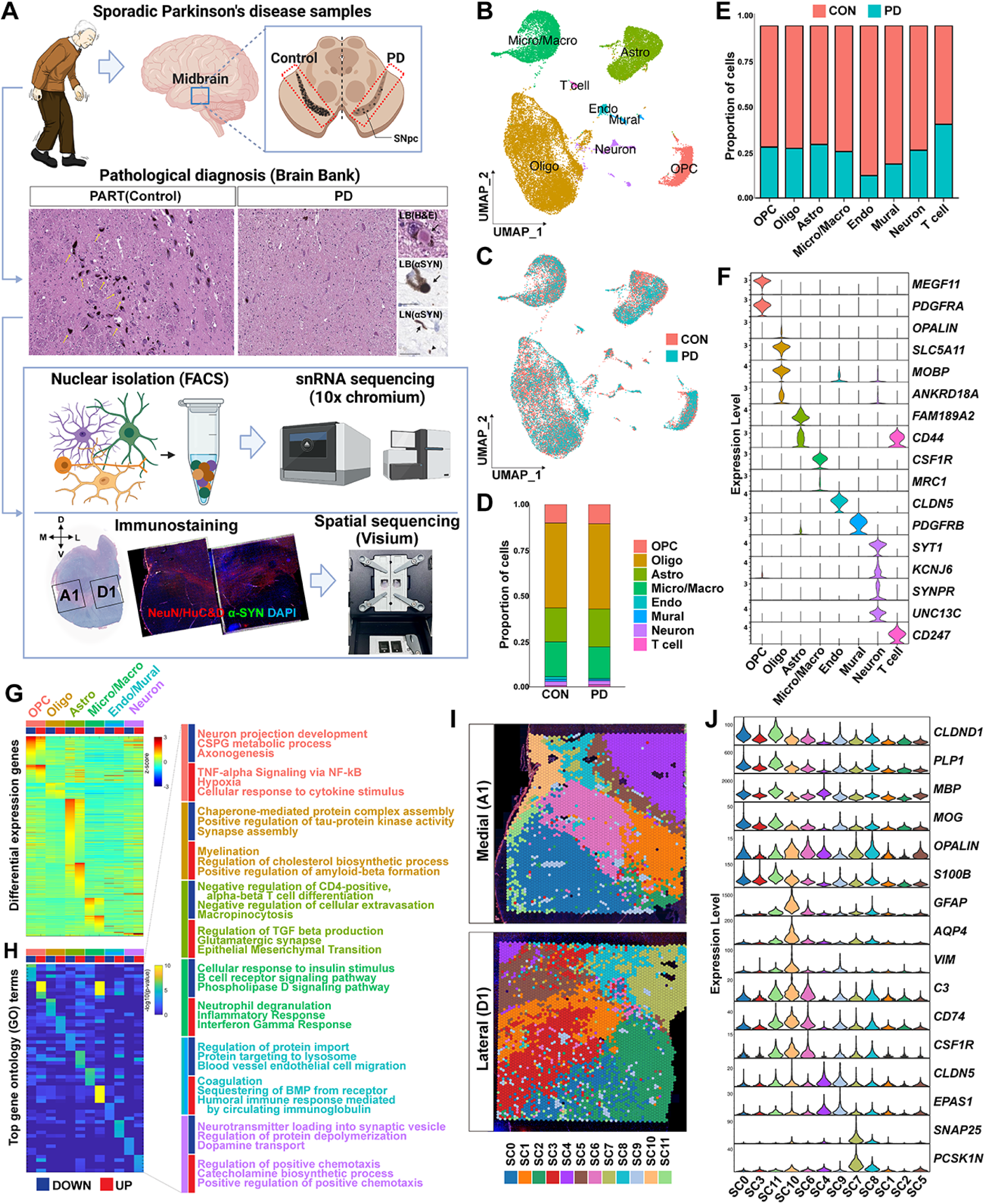
Cell type-dependent expression changes by PD and spatial clustering distribution in human midbrain SN. (A). Schematic of the experimental procedure for snRNA-seq and spatial transcriptomics. Four pathologically confirmed samples each for sporadic definite PD and controls (PART only brains) were used. (B) A UMAP plot of the unsupervised clustering of a total of 33,293 nuclei. (C) A UMAP plot labeled based on the origin of the nuclei. The result shows that CON and PD nuclei are effectively integrated. (D) Proportion of 8 cell types in both Control and PD nuclei; oligodendroglial precursor cell, oligodendrocyte, astrocytes, microglia/macrophage, endothelial cell, mural cell, neuron, and T-cell. (E) Percentage of nuclei from CON and PD samples for each cell type. (F) Selected marker genes for each cell type. (G) Upregulated and downregulated DEGs for each cell type within the PD compared to the control condition. Normalized gene expression scaled for each gene. (H) GO terms for the DEGs identified in (G). The *p*-value is represented in -log10. (I) The Visium spatial clusters (SCs). Each spot within the medial SN corresponding to the A1 capture area and the lateral SN corresponding to the D1 capture area was integrated and categorized into twelve SCs based on their respective gene expression patterns. (J) Representative spatial cluster markers indicating potential cell type composition in each SC.

### Spatial Transcriptomics data analysis pipeline

We obtained two slides of 10x Genomics Visium from two consecutive regions of substantia nigra of post-mortem human brains affected by PD. The datasets were labeled as A22-16_SuN_A1 (A1; Medial) and A22-16_SuN_D1 (D1; Lateral). Scanpy v1.9.8 a Python library (Wolf et al., 2018) was used for spatial data analysis pipeline. We removed spots with mitochondrial gene expression above 20% and more than 35,000 UMI counts. For the A1 data, spots with UMI counts less than 3,000 were removed since most (95%) of those spots were located outside of the tissue. We also removed genes expressed in less than 10 spots. Raw read counts were normalized to the total counts per spots and log-transformed. We performed PCA with HVGs obtained from the Scanpy package. We integrated two spatial slides, Medial and lateral slide, using the Scanorama algorithm [47] to harmonize the datasets. After the integration, we clustered spots using a graph-based algorithm in 100 scanorama embedding space. As a result, we obtained 12 spatial clusters.

### Deconvolution of Spatial Transcriptomics Data

We inferred cell type abundances in the Visium spots using Cell2location, a deconvolution tool [48] based on the cell type expression profile of our snRNAseq data. We filtered out genes expressed in less than 5 spots, 3% of the whole spots, or mean expression less than 1.12 in non-zero spots in the snRNA-seq data. A regression model implemented in Cell2location was instantiated and trained to capture the relationship between cell states and observed gene expression, resulting in the posterior distribution of the estimated cell-type gene expression. We used the intersection of the HVG set in both snRNA-seq and Visium. We then trained the Cell2location model to estimate cell type abundance across spatial locations. This model was extensively trained for 30,000 epochs, using full data batches and GPU acceleration, to ensure robust estimation of spatial cell distribution patterns.

### Trajectory analysis

Monocle2 [49] was applied to reconstruct the transcriptional cellular trajectory from CON to PD. Monocle2 provided a reduced dimension and cell clusters called ‘Monocle components’ using the ‘DDRTree’ method. For each component, we identified DEGs using the same method that was engaged to obtain celltype DEGs. Only genes with an adjusted *p*-value less than 0.05 were considered for further gene ontology and pathway enrichment analysis. To calculate the pseudotime, we decided a root cell cluster as the Monocle component exhibiting the maximum proportion of normal cells. Based on the pseudotime, we manually categorized the Monocle components into several distinct ‘trajectories’’. For each trajectory, we identified pseudotime DEGs using the vector generalized linear model implemented in Monocle2. We used DEGs which have a *q*-value under 0.01 and are expressed in more than 10% of cells in the following analysis.

### RNA velocity

Scvelo [50] were used to predict cellular dynamics of neurons, astrocytes, and microglia along with PD progression. Since neuron subclusters exhibited distinct transcriptional characteristics rather than PD-associated transcriptional changes, we separately applied Scvelo to CON and PD on the same tSNE space. To confirm the changes in the Velocity vectors affected by PD state, we calculated the dot product of the embedded velocity vector of a cell in the CON (or PD) state and the average of the embedded velocity vectors of its three nearest neighboring cells in the PD (or CON) state. For both astrocytes and microglia, we performed Scvelo on the embedded space obtained from Monocle2.

### Gene ontology and pathway enrichment analysis

DEG sets, derived from both Seurat and Monocle2, were queried to an EnrichR [51]. Terms were gathered from either all or a subset of the following databases: ‘KEGG pathway’, ‘GO Biological Process’, ‘GO Molecular Function’, ‘GO Cellular Component’, ‘MSigDB Hallmark’, ‘WikiPathway’, and ‘Reactome’. From the terms with an *p*-value or adjusted *p*-value under 0.05, we selected those with a high combined score, regardless of their origin database.

### GRN analysis

All the neuron subclusters were divided into PD and CON, resulting in 21 groups. NC6 does not have PD cells. Motif-based GRNs were reconstructed for each of the 21 groups using the default SCENIC [52] pipeline. Subsequently, regulon activities within these GRNs were calculated.

For each of trajectories obtained from astrocytes and microglia, we applied TENET [53] a GRN reconstructor based on pseudotime-ordered expression profile. We used the default parameter for the TENET_TF version. We prepared the union of the SCENIC and TENET Transcription Factor (TF) lists, totaling 1708 TFs. To eliminate false positives, we trimmed indirect edges from the generated GRN. We employed Cytoscape [54] to visualize the GRN and merge individual GRNs into a comprehensive network. To identify gene modules, we applied the ‘Leiden Clusterer (remote)’ function of the ‘clusterMaker’ [55] application to the merged GRN with a resolution parameter of 0.65, a beta value of 0.01, and 300 iterations.

### RNAscope and Immunohistochemistry

To minimize variability in the effect of fixation time on in situ results, tissue for RNAscope was prepared separately at autopsy. Midbrain tissue was immediately immersed in 10% formalin and fixed for no longer than 3 days. After preparation of formalin-fixed paraffin-embedded (FFPE) blocks, they were cut at a thickness of 6 micrometers and mounted on coated slides (Fisher scientific #12-550-109). All other procedures followed the manufacturer’s instructions (RNAscope® 2.5 HD Duplex Reagent Kit, ACDBio #AD 322430), including the standard times of 15 min for Target Retrieval and 30 min for Protease Plus treatment, with one minor modification: to improve color development, the time for Hybridize Amp 5,6,9, and 10 was extended by an additional 5 min each. Previously designed Catalog Target probes used for *SLC18A2* (#311431-C2), *TH* (#441651-C1), *GAD2* (#415691-C1), *TCC6* (#1124931-C2), *CHI3L1* (#408121-C2), *CD44* (#311271-C1), *CSF1R* (#310811-C1), and *CD163* (#417061-C2). Slides were scanned with Axio Scan 7 (Zeiss) to acquire whole-slide images, and the resulting images were analyzed with Zen (version 3.4, Zeiss) and QuPath (version 0.5.0) [56].

Immunostaining was performed with the same blocks used for RNAscope. All IHC procedures were performed on the same Ventana automated instruments used for pathology. Briefly, sections were dried overnight in a desiccator, deparaffinized by immersion in xylene, and rehydrated by sequential immersion in a graded series of alcohols. Antigen retrieval pretreatment and peroxidase inhibition were performed to minimize background. A red-colored detection kit (UltraView Universal Alkaline Phosphatase Red Detection Kit, Roche #760-501) was used to avoid confusion of positive staining with pigment neurons in the midbrain. Rabbit Recombinant Monoclonal anti-GAD65/67 antibody (Abcam #ab183999, 1:1500 dilution), Rabbit Polyclonal Tyrosine Hydroxylase (TH) antibody (Abcam #ab112, 1:2000 dilution) were used. The staining was imaged using a slide scanner (Aperio GT 450 DX, Leica Biosystems), and Aperio ImageScope software was used for image processing.

## Results

### Eight different cell types detected in the SN and genes altered by PD

For snRNA-seq analysis, a total of 33,293 nuclei were extracted from the SN of a total of 8 donors, with 4 donors from the CON group and 4 from the PD group, followed by FACS, and snRNA-seq procedure of the 10x Chromium platform (Fig. 1A and Supplementary Table 1). Visium spatial gene expression study was performed using two capture areas, namely the “medial” (A1) and “lateral” (D1) regions, from one midbrain FFPE section of a PD sample (Fig. 1A, sFig. 1A and Supplementary Table 1). Immunofluorescence image stained for phosphorylated α-Syn (green), neuronal marker (HuC/D and NeuN, red), and DAPI, was aligned with the capture areas to confirm the spatial locations of the Visium spots. The distribution of TH and Calretinin immuno-reactivity around the SN was also verified in adjacent sections (sFig. 1B).

First, in snRNA-seq data, eight different cell types formed clusters (Fig. 1B), and the integrated CON and PD samples overlapped well on the UMAP 2D space (Fig. 1C). There were no significant differences in proportions of cell types between CON and PD (Fig. 1D). However, fewer cells were obtained from the PD tissue compared to the CON tissue despite using a similar amount of tissue (total CON nuclei = 22631, total PD nuclei = 9129), and this pattern was consistently observed across all cell types (Fig. 1E).

The predominant cell types observed were oligodendrocytes, astrocytes, microglia and oligodendroglial precursor cells (OPCs). Additionally, a small population of *MRC1*-positive brain-resident macrophages was identified within a microglia cluster (Fig. 1B, F, Supplementary Table 2_1) [57,58]. The neurons were organized into several distinct small clusters, and a cluster with characteristics of T cells was also detected, expressing high levels of *CD44*, known to be involved in cell trafficking into the central nervous system (CNS) [59,60]. Vascular cells were clearly divided into clusters of endothelial cells and mural cells including pericytes, based on known markers such as *CLDN5* and *PDGFRB* [61,62].

We examined the DEGs upregulated or downregulated in PD compared to CON for each cell type (Fig. 1G and Supplementary Table 2_2). We found that the enriched gene ontology (GO) terms and KEGG pathways were well separated by both cell type and PD, indicating cell type-specific functions that may be compromised in disease circumstance or cell type-specific cellular responses (Fig. 1H). For example, downregulated genes in OPCs were linked to processes such as neuron projection development, neuron maturation, chondroitin sulfate proteoglycan metabolism contributing as a major component of the extracellular matrix (ECM), or axonogenesis-related genes implying axonopathy [63–65]. On the other hand, upregulated genes were associated with neurodegenerative environmental factors, including hypoxia and inflammatory cytokines.

We next investigated whether these transcriptomic signatures observed in our data were also associated with genetic risks previously studied in PD patients. Of the DEGs between CON and PD for each of the cell types in Fig. 1G, we compiled a list of PD-associated genes that had been identified in GWASs (including Korean GWAS), multi-omics, and genetic databases to see if they showed differential expression in our data [14,16,20,40–46]. We found that many of the DEGs were associated with specific cells, with changes in OPCs, astrocytes and microglia being particularly prominent. This confirms that the DEGs we found recapitulate changes that are consistent and common in PD, particularly in glial cells (sFig. 2 and Supplementary Table 2_3).

**Figure 2.**
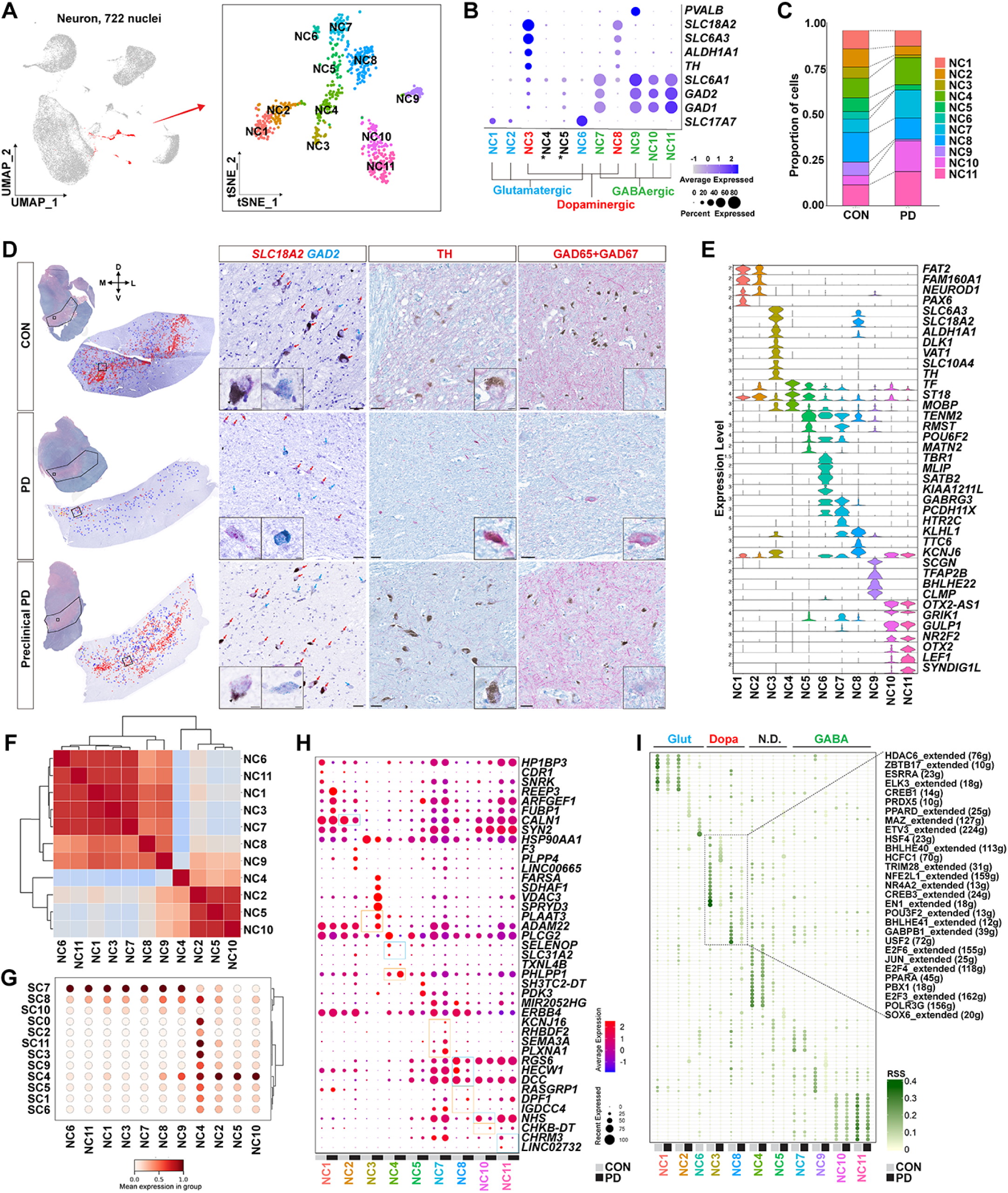
Neuronal subpopulations and changes detected in snRNA-seq and spatial transcriptomes in response to PD. (A) Neuronal subclustering yielded 11 distinct subtypes. tSNE coordinate was used to represent the neuronal subclusters (NCs). (B) Neurotransmitter-based neuronal classification. The asterisk (*) indicates the two clusters that were not specifically characterized. (C) Proportional divergence of NCs between CON and PD. (D) Validation of the neuronal population changes using RNAscope in SN sections from CON (top panel), PD (middle panel), and preclinical PD (bottom panel). The first column shows SN location (polygon) and orientation confirmed in the LFP-stained adjacent section, and each mRNA probe positive cells were manually marked in the enlarged SN image. The second column is an enlarged image of a portion of the SN (small square) in the first column. Scale bar, 50 μm. Cells containing *SLC18A2* (red) and *GAD2* (blue) mRNA strands are indicated by arrows. The third and fourth columns show immunostaining for TH and GAD65+GAD67, respectively, in another adjacent section. Scale bar, 50 μm. Insets show an image of a representative cell. Scale bar, 10 μm. (E) Selected marker genes for each neuronal subtype. (F) Pairwise correlation between NCs based on Cell2location results resolved into spatial transcriptomes. The data show the combined result of the medial and lateral SN capture areas. (G) Scaled mean abundance scores of NCs for each SC. (H) DEGs between CON and PD in each NC. In particular, genes that show statistical significance in both RNA expression and velocity were highlighted with boxes. Blue boxes indicate genes with higher expression in CON, while orange boxes indicate genes with higher expression in PD. (I) Regulons identified in CON and PD for each NC as determined by SCENIC analysis. The color intensity of each dot represents the Regulon Specificity Score (RSS), while the size of the dot represents the *z*-score. N.D. indicates NC4 and NC5 whose properties were not determined.

To elucidate the spatial distribution of cell types in the SN of PD patients, unsupervised clustering on Visium spots resulted in 12 clusters (Fig. 1I and sFig. 3). Despite being initially clustered without spatial information, the Visium spots showed a clear spatial pattern that distinguished the SN and surrounding midbrain anatomical features. The top-ranked spatial DEGs included several cell type markers, allowing each cluster to be distinguished by enriched cell types (Fig. 1J, sFig. 3C, Supplementary Table 3).

**Figure 3.**
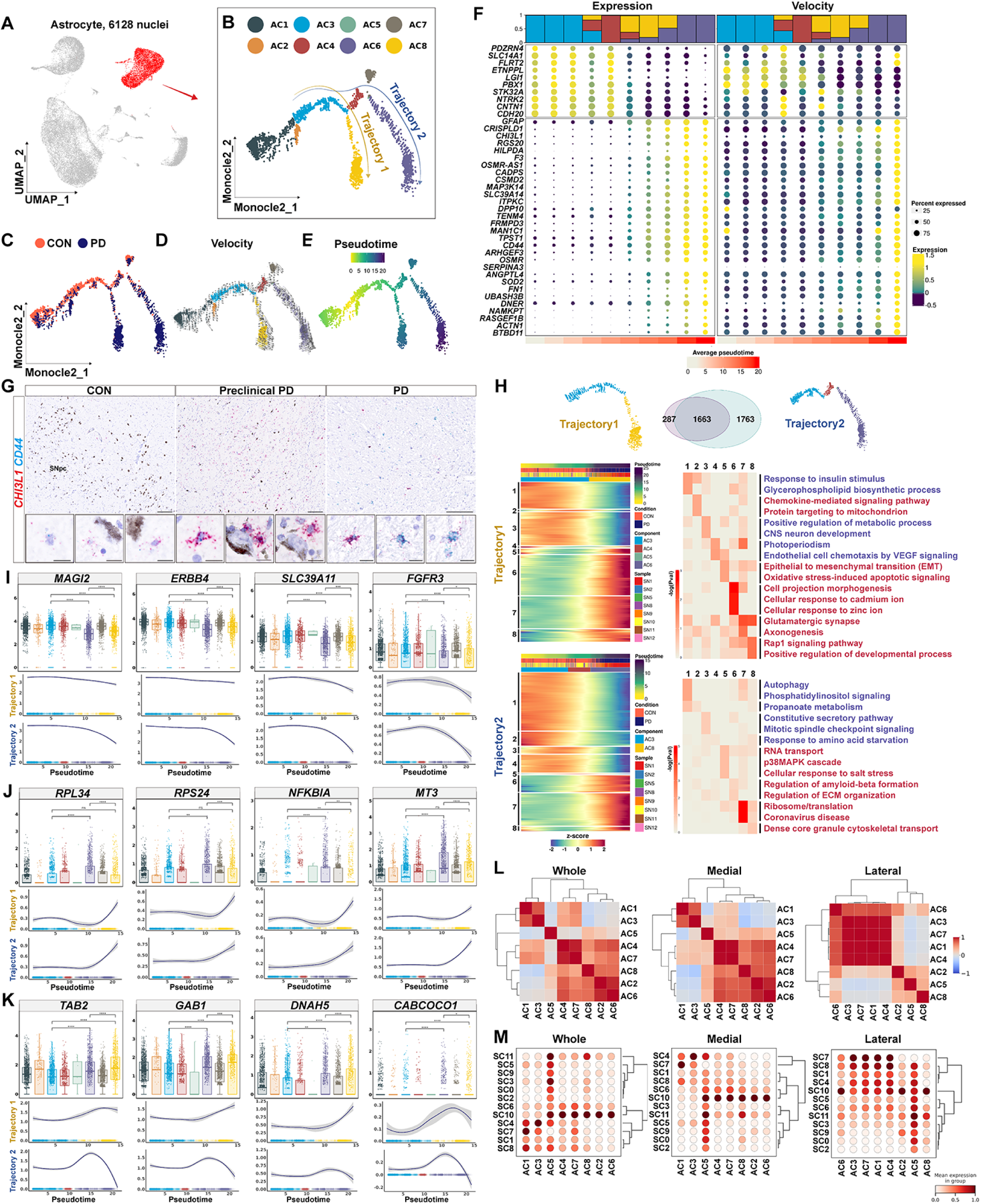
PD-associated pseudotime trajectory analysis in astrocytes and spatial deconvolution of identified subclusters. (A, B) Monocle analysis in the astrocytes subset yielded eight distinct components. Two distinct astrocyte trajectories were indicated. (C, D, E) Monocle coordinates show the distribution of cells divided by the origin of the cells, the magnitude of the RNA velocity vector, and the calculated pseudotime, respectively. (F) Progressive expression (left panel) and velocity (right panel) changes of DEGs on the 10 separate pseudotime slots. Above the plots is the proportion of Monocle components corresponding to each pseudotime slot. Below the plots is the average pseudotime of each slot. (G) Validation of increased expression of DEGs using RNAscope in SN tissues from CON (left panel), preclinical PD (middle panel), PD (right panel). Preclinical PD shows a remarkable increase in *CHI3L1*-positive (red) cells, while definite PD shows prominent *CD44*-positive (blue) cells. (H) Expression changes of Monocle DEGs along trajectory 1 (top panel) and trajectory 2 (bottom panel). At the top, the intersection of DEGs of two trajectories was represented in a Venn diagram. DEGs are sorted into different gene sets based on their expression pattern along pseudotime, and then the GO terms of each gene set are shown using another heatmap. The adjusted *p*-value is displayed in *-log10*. (I, J, K) DEGs diminished (I) and increased (J) more in Trajectory 2 rather than Trajectory 1, and showed opposite changes at the end of the trajectories (K). Box plots are normalized expressions grouped by ACs (top panel) and line graphs are locally regressed expression data over each trajectory (bottom panel). Significance of differences, determined by the *Wilcoxon* rank-sum test, is indicated as follows: ****, 0 < *p* < 0.0001; ***, 0.0001 < *p* < 0.001; **, 0.001 < *p* < 0.01; *, 0.01 < *p* < 0.05. *p* indicates an adjusted *p*-value corrected by the *Benjamini-Hochberg* method. ‘ns’ indicates no significant difference. The upper curve corresponds to Trajectory 1, while the lower curve corresponds to Trajectory 2. The gray area around the curve represents the 95% confidence interval calculated using the standard error. (L) Correlations between astrocyte components resolved into spatial data. Left, middle and right heatmap correspond to the whole (medial and lateral combined), medial, lateral SN spatial data. (M) Scaled mean abundance scores of astrocyte components for each spatial cluster. Notably, the heatmap maintains the same region order as the (L) reference.

Specific spatial clusters (SC) 0 and SC3 were identified as oligodendrocyte-rich regions, expected to be white matter. SC10, SC11, and SC6 were glial-rich regions representing progressive gliosis following neuronal loss. SC4 and SC9 are vascular cell-rich regions. SC7 and SC8 were neuron-rich clusters. Among the two neuron-rich clusters, SC7 had more pronounced expression of various neuronal markers than SC8, including *SYT1*, *SLC18A2*, *GAD1*, *GAD2*, *CALB1* and *TAC1* (also known as Substance P) (Fig. 1J and sFig 3C, 4A-D, 4I).

In particular, SC4 was a mixture of different cells, including oligodendrocytes, astrocytes, and vascular cells with a pronounced expression of *PVALB* (sFig. 3C, 4I), exactly representing the known features of the red nucleus (RN). All these results indicate that cell types in the SN of PD patients are organized in a spatially structured manner.

**Figure 4.**
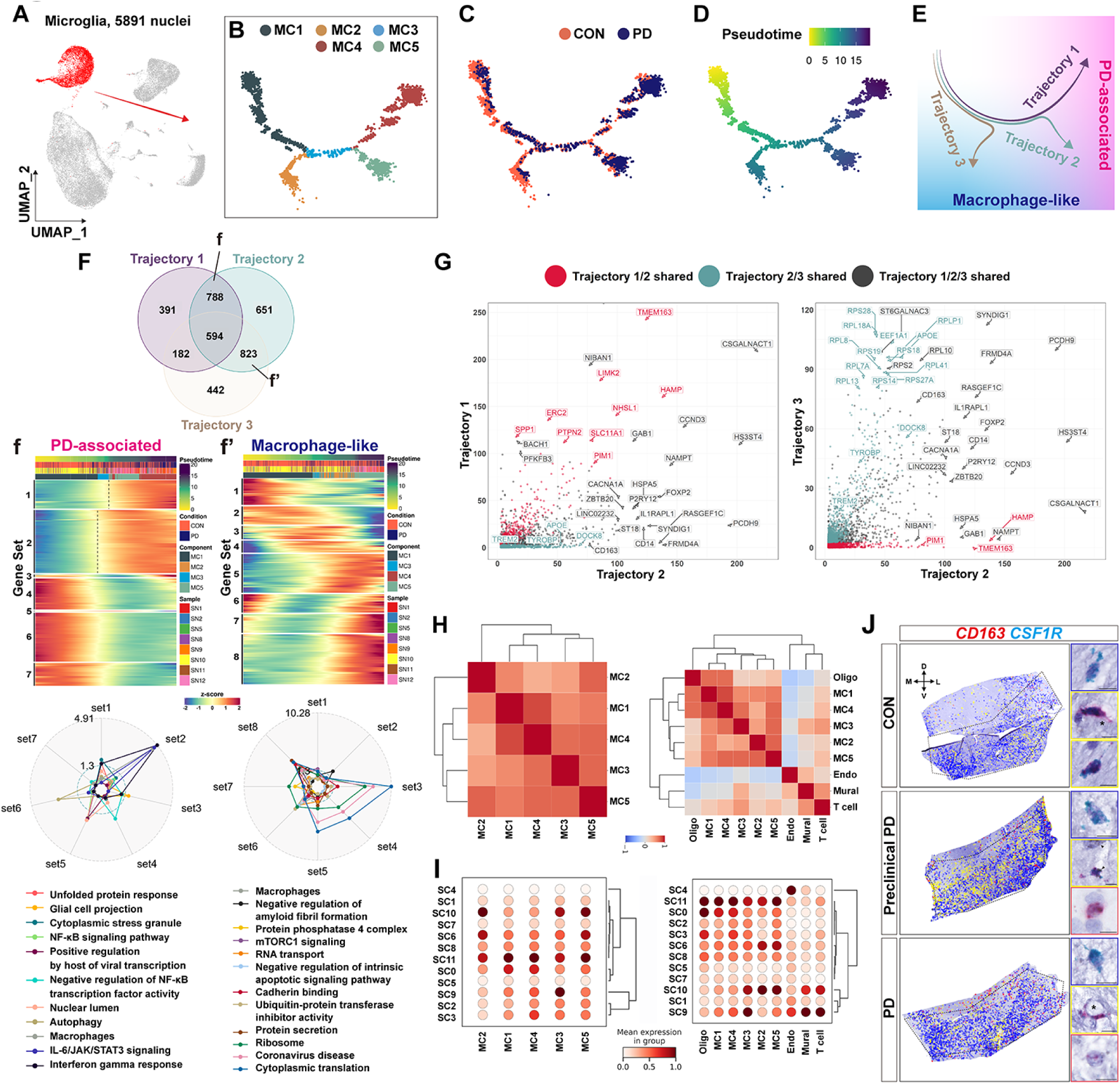
PD-associated pseudotime trajectory analysis in microglia and spatial deconvolution of identified subclusters. (A, B) Monocle analysis in microglial subset yielded 5 distinct components. (C, D) Monocle coordinates showing the distribution of cells divided by the origin of the cells and the calculated pseudotime. (E) Schematic representation of the three identified microglial trajectories and their phenotypes. Trajectories 1 and 2 are increasingly associated with "PD-associated" features (pink gradation), while trajectories 2 and 3 are increasingly associated with "macrophage-like" features (blue gradation). (F) Intersection of DEGs across three trajectories. Heatmaps show the expression of intersecting DEGs between trajectory 1 and trajectory 2 (f) and between trajectory 2 and trajectory 3 (f’). These DEGs are organized as Gene sets according to their expression patterns over pseudotime, with corresponding GO terms shown in the radar plot at the bottom of the heatmaps. The radar plot shows the adjusted *p* value for each term relative to the Gene sets, the *Bonferroni-Hochberg* corrected *p*-value is scaled using *-log10*. (G) Adjusted *p*-value for DEGs relative to each trajectory. The left plot shows the significant genes shared between trajectory 1 and 2 (red), while the right plot shows the genes shared between trajectory 2 and 3 (green). In each plot, the gray dot represents the genes that are common to all trajectories. Each axis refers to the *-log10* scaled *p*-value of specific trajectories, which are *Bonferroni-Hochberg* corrected. (H) Correlation between MCs (left panel), and another heatmap including additional oligodendrocyte, endothelial cell, mural cell, and T-cell features (right panel) resolved into spatial data. (I) Scaled mean abundance scores of MCs (left panel) and another heatmap including additional cell types for each spatial cluster. (J) Validation of expression changes of DEGs using RNAscope in SN tissues from CON (top panel), preclinical PD (middle panel), and PD (bottom panel). Preclinical PD shows a remarkable increase of *CD163* (red) and *CSF1R* (blue) double-positive cells, while definite PD shows prominent *CD163* only positive cells. Insets show images of representative cells for *CSF1R*-only (blue box), double-positive (yellow box), and *CD163*-only (red box) cells in each sample. The asterisk (*) indicates the location of microvessels based on histologic features. It is interesting to note that some of the double-positive cells contain neuromelanin pigments (arrowheads), presumably taken from nearby dopaminergic neurons, which was mainly observed in the preclinical PD section [160].

### Neuronal Heterogeneity in the SN and Population Changes in PD

To observe changes that would be better detected in subpopulations of neurons in the SN, clustering was performed independently on 722 neuronal nuclei, resulting in a total of 11 neuronal subclusters (NC) (Fig. 2A). Examination of expressed neurotransmitters identified NC3 and NC8 as dopaminergic neurons expressing *SLC18A2*, *SLC6A3*, *ALDH1A1* and *TH*. NC7, NC9, NC10, and NC11 were identified as GABAergic neurons predominantly expressing *SLC6A1*, *GAD2*, *GAD1.* NC1, NC2, and NC6 were found as glutamatergic neurons expressing *SLC17A7* (Fig. 2B and Supplementary Table 4).

In a LIGER analysis [39] integrating our data with the transcriptional profile obtained by enrichment of human midbrain neurons recently reported by Kamath et al.[25], we confirmed that the neurons in our data retained the general features of their high-throughput profile of midbrain neuronal subtypes. Specifically, NC3 and NC8 matched well with dopaminergic neuronal subtypes expressing *SOX6* and *CALB1* (sFig. 5).

**Figure 5.**
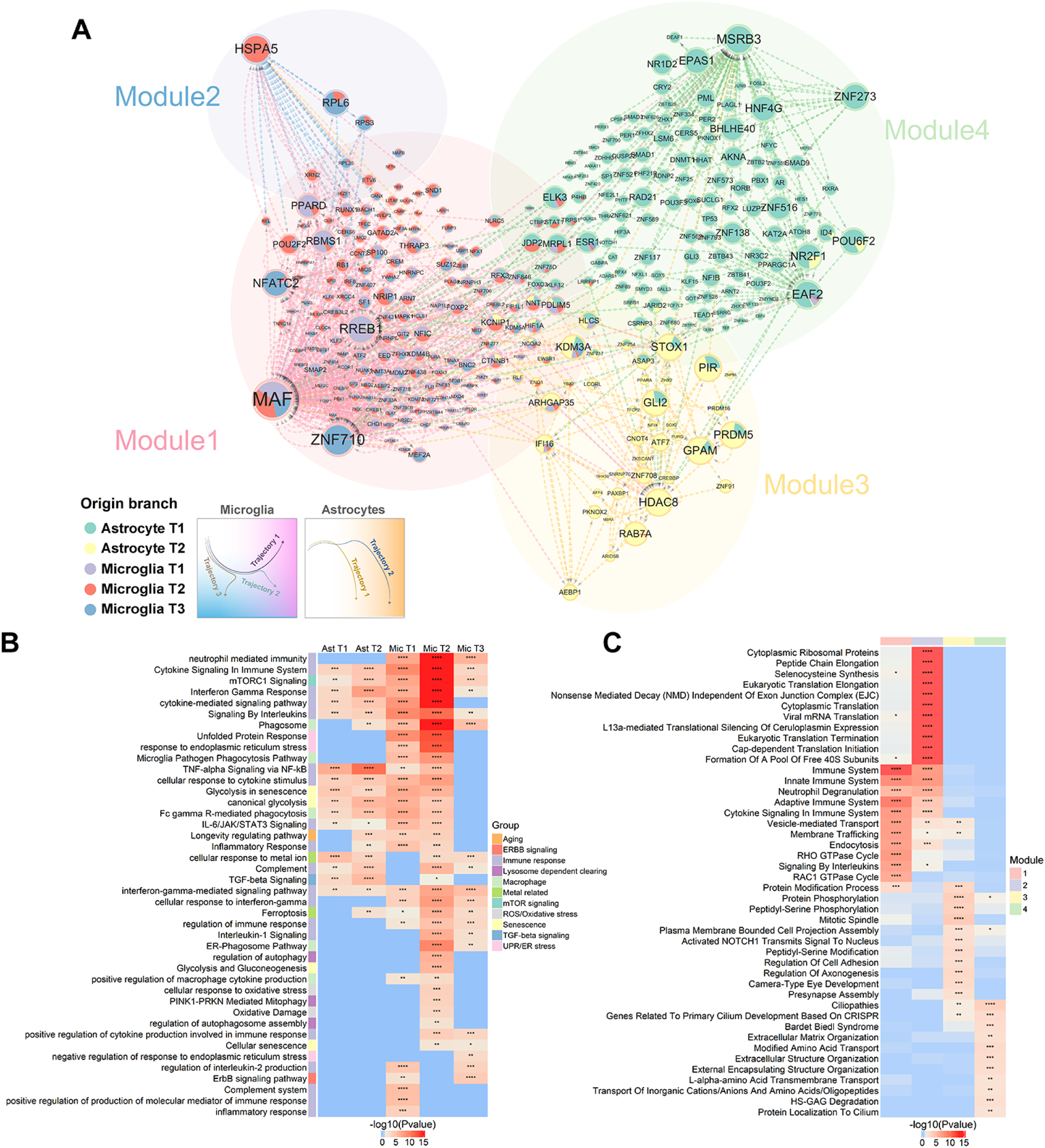
Gene regulatory networks based on TENET analysis using Monocle DEGs from both astrocyte and microglia. (A) Integrated gene networks derived from two astrocyte and three microglia trajectories. Nodes were organized into four distinct modules. The size of each circle corresponds to the number of outdegrees for each node. A pie chart is used to represent the number of outdegrees of each TF prior to the network being merged. (B) GO terms associated with the unique Monocle DEGs for each trajectory. GO terms were grouped by potential PD-associated cellular events or pathways and indicated by color (legend on the right). Significance of terms is indicated as follows: ****, 0 < *p* < 0.0001; ***, 0.0001 < *p* < 0.001; **, 0.001 < *p* < 0.01; *, 0.01 < *p* < 0.05. (C) GO terms associated with the identified modules within the gene network. The networks formed by these modules were not entirely independent as shown in (A), but the transcription factors and their target genes are algorithmically assigned to specific modules, resulting in distinct GO profiles for each module, as shown. The representation of term significance is consistent with that in (B).

By dividing the proportion of identified neuronal subclusters between CON and PD groups in our dataset, we observed a reduction in dopaminergic (NC3 and NC8) and glutamatergic (NC1, NC2 and NC6) neurons, leading to an increased relative proportion of all GABAergic (NC7, NC10, and NC11) neurons, except for the NC9 subcluster, which co-expresses *PVALB* in the context of PD (Fig. 2C) [66].

To examine the proportional changes in neuronal subclusters observed in scRNA-seq result in the tissue, we performed RNAscope using the probes targeting *SLC18A2* as a marker for dopaminergic neurons and *GAD2* as a marker for GABAergic neurons (Fig. 2D). In the CON brain, *SLC18A2* staining was strong in all the pigmented SN neurons while *GAD2*-positive staining was observed in non-pigmented neurons (Fig. 2D, top panel). These GABAergic neurons, known to be predominantly located in the substantia nigra pars reticulata (SNpr) in rodents [67,68], were sporadically scattered and surrounding the SNpc in human samples. On the other hand, in advanced PD, where TH immunoreactivity was significantly reduced, *SLC18A2*-positive neurons had largely disappeared, except for a few remaining in the dorsomedial part of the SN, indicating the loss of dopaminergic neurons (Fig. 2D, middle panel and sFig. 6). *GAD2*-positive neurons were mainly preserved in number relative to *SLC18A2*-positive neurons, indicating the persistence of GABAergic neurons, which may explain the hypokinetic PD phenotypes (Supplementary Table 5) [69,70].

In patients with confirmed synucleinopathy but no manifestation of clinical symptoms, namely preclinical PD, we found that there were still many neurons positive for *SLC18A2* and *GAD2*, similar to those observed in CON. However, the intensity of immunostaining for TH, an enzyme critical for dopamine synthesis, was significantly decreased, which could serve as a good indication of early PD progression (Fig. 2D, bottom panel and sFig. 6).

Upon examining the enriched genes in each neuronal cluster, we were able to gain a more detailed understanding of the specific characteristics of neuronal subclusters (Fig. 2E and Supplementary Table 4). For example, among the dopaminergic neurons, NC3, which was highly reduced in number in PD, had high expression of *TH*, *THY1* and *RAB3A*, while NC8, which was relatively less reduced in number in PD, had high expression of *TTC6*, *KLHL1*, and *SEMA6D* (Fig. 2E and sFig. 7). Different groups showed unique gene expressions. Among excitatory glutamatergic neurons, NC1 had high *PAX6* expression, while in inhibitory GABAergic neurons, NC7 had high serotonin receptor 5-HT-2C (*HTR2C*) levels. Some groups, like NC4, shared features with other cell types, showing oligodendrocyte markers such as *TF*, *ST18*, and *MOBP.* NC5 specifically expressed *MATN2,* a gene known to be active in white matter neurons (Fig. 2E) [71].

To investigate the spatial distribution of neuronal subtypes, we deconvoluted the spatial data into neuronal subtype proportions using Cell2location (,sFig. 8A) [48] followed by correlation analysis. This analysis revealed two spatially correlated groups of neuronal subtypes (Figs. 1I-J, 2F, and sFig. 7). The first group, including two glutamatergic subtypes (NC1 and NC6), two dopaminergic subtypes (NC3 and NC8), and three GABAergic subtypes (NC7, NC9, and NC11). The subtypes were mainly located in the region expected to be the dopaminergic nuclei in the midbrain, including particularly two neuron-rich SC7 and SC8 (Fig. 2F-G and sFig. 8).

The second group comprised a glutamatergic subtype (NC2), a GABAergic subtype (NC10), and two unknown neuron types (NC4 and NC5). These subtypes were primarily located in regions outside SN, particularly in SC4 (Fig. 2G and sFig. 8). This spatial separation of the two groups of neuronal subtypes suggests a transcriptomic distinction between the dopaminergic nuclei and other subregions of midbrain neurons, highlighting the unique molecular and spatial characteristics of different neuronal populations within the midbrain.

Next, we explored the genes showing expression differences between CON and PD samples within each neuronal cluster (Fig. 2H, sFig. 9A-D, and Supplementary Table 4). There was a greater number of DEGs between CON and PD within specific neuronal subtypes (sFig. 9A-B) compared to the total neuronal population (Fig. 1G), suggesting subtype-specific molecular changes associated with PD. With the exception of NC6 and NC9, where the cell number was not sufficient to perform DEG analysis due to the significant reduction in cell number caused by PD, we observed DEGs in the remaining clusters with diverse terms. DEGs related to protein processes in endoplasmic reticulum shown in NC7 and microtubule depolymerization in NC1, involved in different cellular processes (sFig. 9A-D) [72–74]. Among the most significant DEGs in each cluster (Fig. 2H), *VDAC3*, known to recruit Parkin and induce mitophagy [75], *PLAAT3*, a lipid-modifying enzyme involved in organelle degradation [76,77], and *ADAM22*, involved in synaptic maturation [78], showed specific increases in dopaminergic NC3. In NC4, genes related to metal ion homeostasis such as *SELENOP*, a selenium transporter known for its protective role in Alzheimer’s disease (AD) and PD [79,80], and the copper transporter *SLC31A2* [81] were downregulated. In NC7, *SEMA3A* and *PLXNA1*, involved in semaphorin signaling and previously implicated in neurodegenerative diseases [82,83], were upregulated. In NC8, specific downregulation of genes such as *RGS6*, which is important for the maintenance of dopaminergic neurons in the ventral SN [84], and *DCC*, which plays a critical role in the development and survival of the ventral midbrain dopamine circuit [85–88], were observed. *HSP90AA1*, known as a stress sensor in neurons, showed a tendency to decrease in multiple clusters affected by PD, including NC2,3,4 and 8 [89,90]. These subcluster-specific changes were consistent with the heterogeneous nature of PD effects on neurons, as indicated by previous ontology results (sFig. 9C, D).

We incorporated RNA velocity analysis to infer the dynamical changes in the neuronal transcriptome (sFig. 9E and Supplementary Table 6). Significant changes in the trajectory of cell development were observed when comparing healthy controls to PD patients. These changes were especially noticeable in the remaining dopamine-producing neurons (NC8). The dot product calculation between CON and PD velocity vectors for NC8 showed significant negative values (one-sided *Wilcoxon* signed-rank test, *p* = 2.64 x 10^-8^), indicating alterations in both unspliced and spliced form of mRNA due to PD (sFig. 9F).

To further understand the regulators of these changes, SCENIC analysis was used to identify regulon activities in each neuronal subtype (Fig. 2I and Supplementary Table 7) [52]. Regulon activities were primarily clustered according to neuronal subtypes, but PD led to specific alterations, particularly in dopaminergic neurons NC3 and NC8 (Fig. 2I, dotted box). In NC3, regulons such as *BHLHE41* and *BHLHE40*, circadian regulators, *NFE2L1*, which is important for dopaminergic neuronal survival, and *TRIM28*, which simultaneously regulates nuclear accumulation of α-Syn and tau, were significantly inactivated in PD [91,92]. In NC8, *PBX1*, involved in protection against oxidative stress and is implicated in midbrain dopaminergic neuronal development and PD, was a specific regulon and significantly inactivated by PD [93]. *SOX6*, a marker gene for the division of SNpc into ventral and dorsal, was highly active in NC8 [94], suggesting distinct populations of dopaminergic neurons with differential susceptibility to PD. Taken together, despite the detected population-level changes in different neuronal subtypes, the transcriptomic changes in the neurons are likely driven by alterations in dopaminergic neurons. This finding highlights the critical role of dopaminergic neurons in the disease progression of PD as a pathological drive.

### Key Transcriptomic Changes Driving PD-associated Astrocyte Trajectories

In astrocytes, we clustered cells with Seurat as in neurons and found that the clusters were not independently separated on UMAP and that transcriptomic features differed significantly between CON and PD rather than between clusters (sFig. 10). Therefore, we performed trajectory analysis on astrocytes to track pseudo-temporal expression changes as the disease progressed. The 6,128 astrocyte nuclei (Fig.3A) were clustered into 8 astrocyte components (ACs) (Fig. 3B and Supplementary Table 8_1), resulted in two distinct astrocyte trajectories, leading to PD-enriched astrocyte components AC8 (Trajectory 1) and AC6 (Trajectory 2) (labels in Fig. 3B, Fig. 3C). Based on RNA velocity prediction [50] and GO enrichment analysis, we hypothesized that both trajectories start at AC3, which expressed genes associated with the most mature functions of normal astrocyte, including calcium signaling, cAMP signaling and long-term potentiation (Fig. 3D-E and sFig. 11) [95–98]. We visualized the expression changes and corresponding RNA velocity of the DEGs identified in the astrocytes cluster shown in Fig. 1G in a large pseudotime stream consisting of AC3-AC4-AC5-AC6-AC8 and observed that they showed a gradual change along a trajectory that was consistent in both analysis methods (Fig. 3F and Supplementary Table 8_1, 9). This also indicates that the pseudotime trajectory exactly represents the PD progression.

Among those genes, we validated expression changes in PD patient tissues, focusing on *CHI3L1* and *CD44*, genes that have been reported to show expression changes associated with the neuronal injury, particularly the former in acute and the latter in chronic responses (Fig. 3G and sFig. 12A) [99–102]. It was observed that *CHI3L1* was significantly upregulated in preclinical PD cases, particularly in SN, and cells displaying strong *CHI3L1* expression were readily adjacent to pigment neurons. Additionally, *CD44* was consistently co-expressed with *CHI3L1* in preclinical PD cases, although its expression did not show a significant increase. Interestingly, in PD cases, we observed the expression of *CD44* exceeded that of *CHI3L1*. These findings indicate that genes exhibiting changes in expression along the astrocyte trajectory could potentially serve as histological markers of reactive astrocytes not only in the end stage but also in the early response.

Next, to explore the differences between the two trajectories, we performed analysis to identify DEGs altered by pseudotime using a likelihood ratio test based on pseudotime spline fitting implemented in Monocle2 and observed transcriptomic changes associated with Trajectory 1 and Trajectory 2 (Fig. 3H and Supplementary Table 8_2). The result showed that 1,663 genes were shared among 1,950 DEGs from Trajectory 1 and 3,426 DEGs from Trajectory 2. Although approximately 85% of DEGs in Trajectory1 shared with Trajectory 2 across the disease trajectory, the specific terms still appeared (Fig. 3H, upper panel, and all gene lists for each Gene set are shown in Supplementary Table 8_3). Trajectory 1 was characterized by increased expression of genes related to metal ion homeostasis (Gene set 6) and glutamatergic excitatory synapses (Gene sets 7-8).

Trajectory 2 had 1,763 DEG genes that do not appear in Trajectory 1 (Fig. 3H, lower panel and Supplementary Table 8_3). This Trajectory 2 exhibited upregulation of genes involved in the p38 MAPK cascade and cellular response to general stress during the AC3 to AC4 transition (Gene set 5), potentially in response to extracellular tau or α-Syn accumulation [103–106]. Also, Trajectory 2 showed unique features, including upregulation of genes related to ribosome biology or coronavirus disease (Gene set 7) at the end state.

DEG analysis showed downregulated genes in AC6 compared to AC8, including *MAGI2, CADM2* and *NEBL* (cell-cell interaction molecules)*, ERBB4,* and *SLC39A11* (metal ion transporter) (Fig. 3I and sFig. 12B). Genes upregulated in AC6 compared to AC8 included several ribosomal proteins (*RPL34*, *RPS24*, *RPL41*, *RPS23,* and *RPS25*) (Fig. 3J and sFig. 12C). Genes that are highly expressed in AC8 opposed to AC6 included cilia-related genes (*DNAH5* and *CABCOCO1*) and the subunits of mitochondrial ATP synthase (*ATP5F1E* and *ATP5ME*) (Fig. 3K and sFig. 12C). In addition, the difference in *FGFR3*, known as a suppressor of *GFAP* expression along with astrocyte reactivity [107,108], as well as the alteration in *NFKBIA* and *TAB2*, the NF-kb activator, suggest the major signaling pathways may involved (Fig. 3I-K).

Taken together, the two trajectories of astrocytes, although sharing many similarities, underwent specific genetic changes at the end, suggesting the presence of different types of reactivity in astrocytes or different circumstances in the damaged area.

Spatial correlation analysis revealed that astrocyte subtypes were primarily grouped into two categories based on their association with PD (Fig. 3L-M and sFig. 8B): CON-like (AC1 and AC3) and PD-associated (AC4, AC7, AC8, AC2, and AC6) (Fig. 3L, left panel), which are actually aligned with their pseudotime stages (Fig. 3B). The CON-like astrocytes were located in neuron-rich regions (SC7 and SC8) and the *PVALB*-high SC4 region. PD-associated subtypes were located in the glia-rich regions (SC10, SC11 and SC6), indicating ongoing reactive gliosis (Fig. 3M, left panel and Supplementary Table 3). Interestingly, in regions with PD-associated subtypes, there was a clear scarcity of CON-like astrocytes, indicating a mutually exclusive population distribution and possibly representing different states of the same cells.

Spatial correlation between late-stage subtypes AC6 and AC8 was high in the medial SN but low in the lateral SN, resulting in completely different groupings in hierarchical clustering (Fig. 3L). In this regard, the abundance score of AC6 was higher than AC8 in SC7 and SC8, but lower in SC0 and SC2, with this difference being more pronounced in the lateral SN (Fig. 3M, right panel). In the medial SN, AC4 and AC7, showed high spatial correlation with PD-derived astrocytes (AC8, AC6 and AC2), but in the lateral SN, these showed high correlation with CON-like astrocyte (AC3 and AC1) (Fig. 3M, middle and right panels). These results indicate the spatiotemporally different effects of PD exerted on medial and later SN [109,110], which may be related to the different proportion and distribution of AC6 and AC8 astrocyte subtypes in these regions.

### Identification of two distinct character transitions in PD microglia

5,391 microglia nuclei were sorted by pseudotime trajectory, displaying five microglia components (MCs) composed of one trunk and four symmetrical branches (Fig. 4A-B and Supplementary Table 10_1). Like astrocytes, microglia trajectory was also separated by CON and PD (Fig. 4C). Moreover, according to the hierarchical enrichment GO analysis, the MC1 corresponds to normal microglia, which regulate homeostasis through AMPK signaling and microtubule anchoring (sFig. 13). Three trajectories were identified: MC1-MC3-MC4 (Trajectory 1), MC1-MC3-MC5 (Trajectory 2), and MC1-MC2 (Trajectory 3) (Fig. 4D-E). Trajectory 1 and 2 (Fig. 4C and Supplementary Table 10_2) shared 788 pseudotime DEGs were categorized into seven Gene sets (Fig. 4F, 4f, and all gene lists for each Gene set are shown in Supplementary Table 10_3). Gene set 2 was associated with interferon-gamma response, macrophages, and IL-6/JAK/STAT3 signaling, indicating inflammatory response. The subsequent upregulation of Gene set 1 in MC5 suggested a full-blown stress response, including UPR and stress granules. A small Gene set 5, whose expression is transiently increased before and after MC3, was associated with dynamic changes in the nuclear lumen and the NF-kB signaling pathway. The largest Gene set 6 showed downregulation of autophagy-related genes.

Next, we noted the common features of downward trajectories 2 and 3. They shared 823 pseudotime DEGs divided into eight gene sets (Fig. 4F, 4f’, and Supplementary Table 10_3). Gene sets 3, 4, and 5 were associated with ribosome and related terms, cytoplasmic translation and coronavirus disease. Gene sets 3 and 4 showed a pattern of transiently increased expression in the middle of the pseudotime, followed by Gene set 5 including the additional term, protein secretion. There was a rebound in the genes of Gene set 6 was associated with ubiquitin protein transferase inhibitor activity, negative regulation of intrinsic apoptotic pathways, and mTORC1 signaling. Gene sets 7 and 8 were upregulated at the end of the pseudotime and were associated with microglial activation, maturation, immune function, and protein secretion.

These results suggest the distinct microglia trajectories associated with PD progression, highlighting the critical role of the MC3 state in determining the fate of microglia and the involvement of inflammatory responses, stress responses, and dysregulated cellular processes in disease-associated trajectories.

We then utilized the *q*-value of pseudotime DEGs to identify key genes for two phenotypes (Fig. 4G and Supplementary Table 10_2): one for the “Macrophage-like” phenotype (shared in Trajectory 2 and 3) and the other for the “PD-associated” phenotype (shared in Trajectory 1 and 2) (as labeled in Fig. 4E). The former phenotype was enriched with genes involved in ribosomal biogenesis or protein translation, such as *RPS28*, *RPL18A*, *RPS18*, *RPS19*, and *RPLP1*, consistent with our GO results (Fig. 4G, right panel). *EEF1A1* (microglial polarization) [111] and macrophage markers *TYROBP* [112] were included. *APOE* [113], *TREM2* [114–116] and *DOCK8* [117], which have been implicated in phagocytic function, microglial activation, and relevance to neurodegenerative diseases, were also included. The later phenotype was enriched with genes linked to metal ion transport or ferroptosis such as *TMEM163*, *SLC11A1*, and *HAMP*, which showed high *q*-values [118–122] (Fig. 4G, left panel). Other top genes shared in all trajectory include *CSGALNACT1* (an enzyme for the synthesis of chondroitin sulfate proteoglycan and major component of the glial scar) [123], *NIBAN1* (stress response) [124], *GAB1* (M2 polarization) [125,126], and *CD163* (macrophage marker) [127]. It also has *BACH1* (defense against oxidative stress, mitochondrial dysfunction, and neuroinflammation) [128,129], and *HSPA5* (known as GPR78 or BIP, a regulator of UPR and ER stress) [130,131].

To examine the spatial distribution of microglia, we examined the pairwise correlation between the abundances of five microglial components in the two Visium slides (Fig. 4H, left panel). Unlike neurons and astrocytes, the five microglial components were highly correlated indicating a similar distribution, which was more pronounced in lateral SN (sFig. 8C). MC1 and MC4 associated with Trajectory 1 were spatially correlated with oligodendrocytes (Fig. 4H, right panel). Consistent with this, MC1 and MC4 were enriched in the oligodendrocyte-rich regions SC11 and SC0 (Fig. 1J, 4I). In contrast, SC10 showed enrichment of the "macrophage-like" MC2, MC3 and MC5 rather than MC1 and MC4 (Fig. 4I). In addition, given that MC2 (Trajectory 3) had PD-independent features in snRNA-seq (sFig. 14C), MC5 was relatively enriched in SC9 (perivascular), SC2 and SC3 compared to MC2 (Fig. 1I). SC9 also showed high MC3 abundance.SC4 and SC5 had significantly fewer microglia.

We sought to confirm these analysis results in postmortem brain tissues using RNAscope (Fig. 4J and Supplementary Table 5). *CSF1R* is a well-known marker gene predominantly expressed in CNS microglia (Fig. 1F). In the snRNA-seq analysis, *CSF1R* showed significantly decreased expression specifically in the end of the Trajectory 2, which was associated with the “PD-associated” microglial phenotype (Supplementary Table 10_2). *CD163*, known as a marker of macrophages and previously associated with the anti-inflammatory M2 type, showed increased expression in MC2, MC3 and MC5, associated with the "macrophage-like" phenotype. However, *CD163* relevance was negligible in MC4, which was the end product of Trajectory 1 (“PD-associated” phenotype) (sFig. 15A and Supplementary Table 10_2). In CON tissue, approximately 17% of microglia co-expressed *CSF1R* and *CD163* with perivascular localization (Fig. 4J, top panel). In preclinical PD, the density of *CSF1R*-positive cells increased, with an increase in *CSF1R*/*CD163* double-positive cells (around 36%) (Fig. 4J, middle panel), indicating early and rapid contribution of *CD163* (potentially MC3) (sFig. 15 and Supplementary Table 10_1). In advanced PD, the number of cells positive for *CD163*, but lacking *CSF1R* was markedly increased, consistent with the snRNA-seq results.

### Identification of four distinct Gene Regulatory Network (GRN) modules in PD glial cells

We constructed the GRN between transcription factors (TFs) and DEGs across the identified trajectories/phenotypes in astrocytes and microglia using TENET [53] and Leiden clustering [132] (Fig. 5A, 5B, Supplementary Table 11). Four distinct GRN modules were identified (Fig. 5A, C): Module 1 was a large module comprising all three microglial trajectories, enriched for immune system, cytokine secretion, and vesicle transport processes. Module 2 was a small module, centered on *HSPA5*, linked to UPR and ribosome biology changes in PD microglia. Module 3, associated with astrocyte trajectory 2, was enriched for mitosis, cell adhesion, potentially tau phosphorylation. Module 4 was associated with astrocyte trajectory 1, enriched for cilia-related function, primarily regulated by the TFs, *MSRB3* or *EAF2*.

While the microglial trajectories clustered into one major module (module 1), distinct TFs were found to preferentially regulate specific trajectories within this module. For example, *RREB1* and *PPARD* in Microglial Trajectory 1, *POU2F2* and *XRN2* in Microglial Trajectory 2, and *ZNF710* and *NFATC2* in Microglial Trajectory 3. Considering the prominent role of *HSPA5* in module 2 (blue background circles, Fig. 5A) and its association with the UPR [133,134], the diverse translation-related terms in GO suggest that changes in ribosome biology may be a major response of PD microglia to ER stress, presumably detrimental considering the PD progression in Microglia Trajectory 2 (Fig. 5C) [135,136].

The astrocyte trajectories formed independent modules (Module 3, 4), suggesting distinct transcriptional regulation (Fig. 5A, C). In Module 3 (Astrocyte trajectory 2, grouped in the yellow circle in Fig. 5A), phosphorylation-related terms are likely the result of the contribution of *STOX1*, which has been implicated in tau protein phosphorylation (Fig. 5C) [137]. On the other hand, Module 4 (Astrocyte Trajectory 1, grouped in the green circle in Fig. 5A), primarily regulated by the transcription factors *MSRB3* or *EAF2*, shows terms specific to cilia-related functions, consistent with the enriched cilia-related genes in AC8 (Fig. 5C, 3K).

Taken together, network analysis integrating TF-gene interactions helped further refine the distinct transcriptional programs and biological roles of the identified glial trajectories/phenotypes in PD pathogenesis.

## Discussion

We performed snRNA-seq on SN from pathologically confirmed PD cases and PARTs from SNUH brain bank for the first time. Using PART tissue as controls minimized heterogeneity within the comparison groups, allowing better detection of PD-associated changes [28,29]. The scRNA-seq analysis distinguished 11 neuronal subtypes, including two dopaminergic neuron populations, showing expected population changes and two distinct spatial distributions in PD. Astrocytes and microglia each showed two distinct trajectories/phenotypes associated with PD progression based on DEG.

We validated some of the PD-associated expression changes (e.g., *CHI3L1* in astrocytes, *CD163* in microglia) in the midbrain tissues across PART, preclinical PD, and definite PD cases using the RNAscope assay. Notably, the preclinical PD case showed more pronounced expression of reactive astrocyte marker *CHI3L1* and “macrophage-like” microglia marker *CD163* compared to the late stage PD, suggesting snRNA-seq data captures early transcriptomic changes. By integrating snRNA-seq data with spatial transcriptomics, we gained a comprehensive understanding of how PD affects different cell populations and their spatial distributions in dopaminergic midbrain nuclei and its surrounding regions.

Previous single-cell transcriptomic studies have shown varying results, with some observing little change even in dopaminergic neuron numbers, while others found a specific decrease in a subpopulation of dopaminergic neurons in PD, with no significant changes in other neuronal subtypes [23,24,26]. In the current study, with carefully selecting samples and integrating with histological and spatial information, the expected changes in the SN neurons were accurately recapitulated. A decrease in dopaminergic neurons and an overall reduction in the glutamatergic populations were observed, with the remaining neurons being *GAD2*+ GABAergic neurons (Fig. 2C). Of the two dopaminergic clusters (NC3 and NC8), NC3 appeared more susceptible, exhibiting different gene expression patterns (sFig. 7) and spatial distribution than NC8 (Fig. 2F,G). However, NC3 did not express previously identified markers of vulnerable dopaminergic populations like *SOX6* nor *AGTR1* [24,25]. The remaining dopaminergic neurons in the dorsomedial SN of the definite PD tissue (Fig. 2D, middle panel) were speculated to be the "survived" NC8 population, which was confirmed by examining the location of *TTC6*-expressing neurons using RNAscope (sFig. 16). We cautiously assert the *TTC6* as a potential marker of resistant population of dopaminergic neurons, but further research is needed to understand their distribution in the control and why they are less vulnerable.

The GABAergic neuron is primarily involved in inhibitory functions in the SNpr and is known to be heterogeneous [138]. This study identified four GABAergic subpopulations in the SN region using snRNA-seq. One of these subpopulations, NC9, expressed *PLVAB* and appeared particularly susceptible to degeneration in PD, one of the GABAergic subtypes previously reported to exist in the lateral SNpr [138,139]. The remaining three GABAergic clusters were relatively protected from PD-associated degeneration, as confirmed by RNAscope with the *GAD2* probe. Unlike the dopaminergic neurons which clustered in the dorsomedial SN (Fig. 2D, middle panel), the GABAergic regions had a more scattered distribution confirmed by GAD2 (GABA synthesis enzyme) transcripts using RNAScope. Basically, the boundary between SNpc containing dopaminergic neurons and SNpr containing GABAergic neurons appeared less distinct in humans, compared to rodents as known (Fig. 2D, top panel). These findings support the idea that loss of dopaminergic neurons is the primary pathology in PD, consistent with the SCENIC result (Fig. 2I), and add insight into the recent interest in the GABAergic pathway associated with hypokinesia and other accompanying symptoms of PD [69,140]

The integration analysis of spatial transcriptomes of our data revealed two distinct major groups of neuronal clusters with distinct spatial distributions in the SN and VTA regions: NC6,11,1,3,7,8,9 and NC2,5,10 (Fig. 2F). Three neuron-abundant spatial clusters (SC7, SC8, and SC10) were identified as corresponding to the VTA based on their anatomical locations (Fig. 2G). The SC7 was the most neuron-rich area with low microglial presence and high abundance of control (non-reactive) astrocytes (AC1 and AC3), suggesting it was relatively intact (Fig. 2G, 3M, 4I). In contrast the neighboring SC8 region exhibited lower neuronal abundance and higher association with glia like astrocytes and microglia, indicating it may be more affected by PD pathology. The SC10 region, expected to be ventromedial VTA (vmVTA), exhibited even lower neuronal clusters compared to SC8. It exhibited a high abundance of astrocyte-subtypes, associated with PD progression (AC5, AC4 and AC7) as well as highly PD reactive phenotypes (AC2, AC6 and AC8). It also showed high abundance of microglia (MC3), associated with either progressing to PD or macrophage-like populations (MC2 and MC5) associated with the active PD progression. The findings suggest that the transcriptomic alterations observed in astrocytes and microglia during the progression of PD in the SN region are comparable to those occurring in the VTA. However, despite these parallel glial responses across the two midbrain regions, our data confirmed the well-established notion that neurons in the VTA exhibited reduced susceptibility to degeneration when compared to the vulnerability of neurons in the SN [141–143].

SC4 is thought to be the RN of the midbrain, while the surrounding SC5 region is considered the parabrachial pigmented nucleus (PBP) in the VTA. The neuronal clusters NC2, NC5, and NC10 showed high correlation with SC4/RN, consistent with containing sparse non-dopaminergic neurons. However, since efforts were made to avoid the RN during sample preparation for snRNA-seq, the detection of the SN neuronal subtypes in SC4/RN is specific to the PD case used, potentially due to the increased RN size observed in PD [144]. SC6 and SC1 (expected to be SNpc) clustered with SC5 and SC4 (Fig. 2G), suggesting some similarity but little redundancy of PD-associated reactive astrocytes (AC8 and AC6) or microglia (Fig. 3M, 4I) in this region, which are not directly involved in PD pathogenesis. The presence of neuronal clusters NC8 (sFig. 16B), and NC9 in SC4/RN (sFig. 9I) could be explained by detection of their marker genes (*TTC6* and *PVALB*) in the RN [145,146]. Importantly, SC4/RN showed a very high abundance of endothelial cells, consistent with the RN being a highly vascularized structure (Fig. 4I, right panel) [147,148].

We performed a pseudotime-based trajectory analysis on astrocytes and microglia using Monocle to focus on the transcriptomic changes with PD progression. The analysis aligned individual cells based on their transcriptomic features, regardless of their origin, CON or PD cases, accounting for potential ‘control-like cells’ in PD samples [149]. The distribution of PD-origin cells increased along the pseudotime trajectories, and transcriptomic changes were associated with various cellular events occurring during PD progression. In microglia, inflammation-related processes like interferon-gamma response, IL-6/JAK/STAT2 signaling, and phagocytosis increased along the trajectory (Fig. 4f, gene set 2), while autophagy genes decreased, suggesting inflammation as an initial driver (Fig. 4f, gene set 6) [150,151]. Furthermore, the UPR and cytoplasmic stress granule genes were subsequently increased (Fig. 4f, gene set 1), indicating a sequential induction of the pathogenic process.

Microglia Trajectory 3 showed no influence of inflammation and minimal association with PD. It shared 823 monoclonal DEGs with Trajectory 2, but most cells ended up in the CON fate (Fig. 4C, 4E). GO analysis of unique DEGs revealed their involvement in the lysosomal pathway process, including primary lysosomes, aggrephagy, lysosomes, and autolysosomes (sFig. 14, top panel). This enrichment of the lysosomal pathway may indicate increased lysosomal function with aging [152,153] and "primed" microglia susceptibility [154,155].

This idea can be supported by the observation of spatial transcriptomics. MC3 is more spatially correlated with MC2-MC5 than with MC1-MC4. MC2 has a similar spatial abundance pattern to MC5 and MC3, suggesting a state transition under the influence of PD in the same region (Fig. 4I). MC1 also had a high spatial correlation and similar abundance pattern to MC4, suggesting the same type of microglia in a different state. The higher abundance of MC2-MC5 rather than MC1-MC4 in regions SC6 and SC10 is consistent with the observed active PD progression. The higher abundance of MC5 compared to MC2 in SC9, SC2 and SC3 indicates where PD has already attacked. Given the large number of downregulated genes during the transition from MC1 to MC4 (Trajectory 1), this is more likely a passive or chronic population.

Microglia are commonly divided into two phenotypes, M1 and M2, which have opposing roles in disease [156]. We examined whether the bifurcated Monocle trajectories identified in the same PD sample corresponded to the classical M1 and M2 phenotypes. We analyzed expression changes of known M1 markers, *MRC1* and *CD163,* and M2 markers, *FCGR2A* (CD32), *FCGR1A* (CD64), *PTPRC* (CD45) and *CD14* along the pseudotime trajectories (sFig. 15B, 15A, respectively). Neither trajectory corresponded perfectly to a specific polarization phenotype, suggesting they do not simply represent these classical phenotypes [156–158].

We also examined the disease-associated microglia (DAM) markers, identified in AD research using scRNA-seq [116], in our data set (sFig. 15C). Genes upregulated in DAM like *CLEC7A*, *TREM2*, and *APOE* were involved in both MC2 and MC5 trajectories. However, some DAM markerslike *CD9,* and *GPNMB* associated more with MC2/Trajectory 3, less relevant to PD compared to MC5. This discrepancy could be due to differences and variations across neurodegenerative diseases. Nevertheless, we believe that the characteristics of the microglia trajectories we identified can help distinguish between normal microglia activation and PD-related activation, among known reactive microglia markers. Therefore, we consider the *NLRC5* gene among DAM as a promising PD-associated microglia marker, as its expression increases in the inflammation-related MC3 and continues in MC4 and MC5, but remains low in MC2 (sFig. 15, red boxes). Another gene showing a similar pattern is *FCGR1A* (CD64), and we believe these two genes may have specific roles in PD-associated microglia, requiring confirmation in further studies.

In astrocytes, two PD-associated trajectories were observed with distinct genetic changes at the end of the pseudotime. However, it was initially unclear what determined the divergence of these trajectories. Genes related to the inflammatory response were observed, but it did not seem to be the early intervention as in microglia. However, GRN analysis revealed they formed different GRN modules, suggesting changes in different cellular functions. One module involved ciliary function (homeostatic), the other protein phosphorylation and the mitotic spindle (reactive state abnormalities). Transcription factors such as *HSPA5*, *HDAC*s, *STOX1*, and *JARID2* implicated in neurodegenerative diseases lend credence to our analysis. It is also a novel finding that *MSRB3*, a gene known for its ability to repair oxidative stress [159], is strongly implicated in the astrocyte trajectory in PD, and that its target genes are associated with cilia. Therefore, it is important to conduct detailed experimental validation to understand whether oxidative stress damages cilia or if proper cilia function helps protect against oxidative stress.

## Conclusions

This study represents the first single-nucleus and spatial transcriptomics analysis in a Korean population expanding our understanding about cell type-specific molecular changes within the SN due to PD. We identified distinct transcriptomic profiles of neuronal subtypes and gained new insights into their differential distribution and PD-induced changes. By elucidating PD-associated glial subtypes and their spatial distribution in midbrain dopaminergic nuclei, the specific landscape of the glia environment induced by dopaminergic neuron alterations in the SN was revealed. Notably, the early involvement of inflammation in microglia and the identification of two functionally distinct GRNs in astrocytes provided significant insights into the progression of PD. These findings are crucial for assessing the impact of drug efficacy being developed by global pharmaceutical companies on East Asia and provides a ground truth for further identification of tailored treatment targets within this population.

## List of abbreviations

AC: astrocyte monocle components
AD: Alzheimer’s disease
CON: Control
DEG: Differentially expressed genes
ER: endoplasmic reticulum
GO: Gene ontology
GRN: Gene regulatory network
MC: microglia monocle components
NC: neuronal subclusters
OPCs: Oligodendrocyte Precursor Cells
PART: Primary Age-Related Tauopathy
PD: Parkinson’s Disease
SC: spatial cluster
SCENIC: single-cell regulatory network inference and clustering
SN: Substantia nigra
SNpc: Substantia nigra, pars compacta
SNpr: Substantia nigra, pars reticulata
snRNA-seq: single-nuclei RNA sequencing
smFISH: single-molecule fluorescence in situ hybridization
TENET: Transfer Entropy-based causal gene
NETwork TF: Transcription factor
tSNE: t-distributed Stochastic Neighbor Embedding
UMAP: Uniform Manifold Approximation and Projection

## Declarations

### Ethics approval and consent to participate

Human brain samples were obtained following protocols authorized by the Institutional Review Board of the Seoul National University Hospital (approval number 1808-087-966).

### Consent for publication

Authors consent for the publication. All authors read and approved the final manuscript.

### Availability of data and materials

The raw sequence data reported in this paper have been deposited in the Korea Sequence Read Archive (KRA) in Korea Bioinformation Center, Korea Research Institute of Bioscience and Biotechnology (KAP240761) and are publicly accessible at https://kbds.re.kr/KRA.

## Acknowledgments

The authors would like to thank the brain donors and their families at the SNUH-BB and the funding sources that made this research possible.

## First Authors information

Three authors contributed equally: Sooyeon Yoo, Ph.D., Kwanghoon Lee, B.S., and Junseo Seo, B.S.

## Contributions

S-H.P. and J-K.W. conceived and designed this study. S.Y. and K.L. acquired the snRNA-seq and Visium spatial data. J.K., S.Y., J.S., H.C., and J-K.W. analyzed and interpreted the data. S.Y. and K.L. performed and analyzed the RNAscope experiment. S-I.K. and S-H.P. provided critical advice on sample selection and overall pathological insights. K.L. Y-M.S, and S.Y. assisted in the initial setup of single nucleus preparation. S.Y., J.K., J.S., and H.C. wrote the manuscript, and S-H.P. and J-K.W. revised it substantially. All authors discussed and commented on the manuscript.

## Emails of the corresponding author

Sung-Hye Park, M.D., Ph.D., Jae-Kyung Won, M.D., Ph.D., and Junil Kim, Ph.D.

## Supplementary Figure legend

**Supplementary Figure 1.**
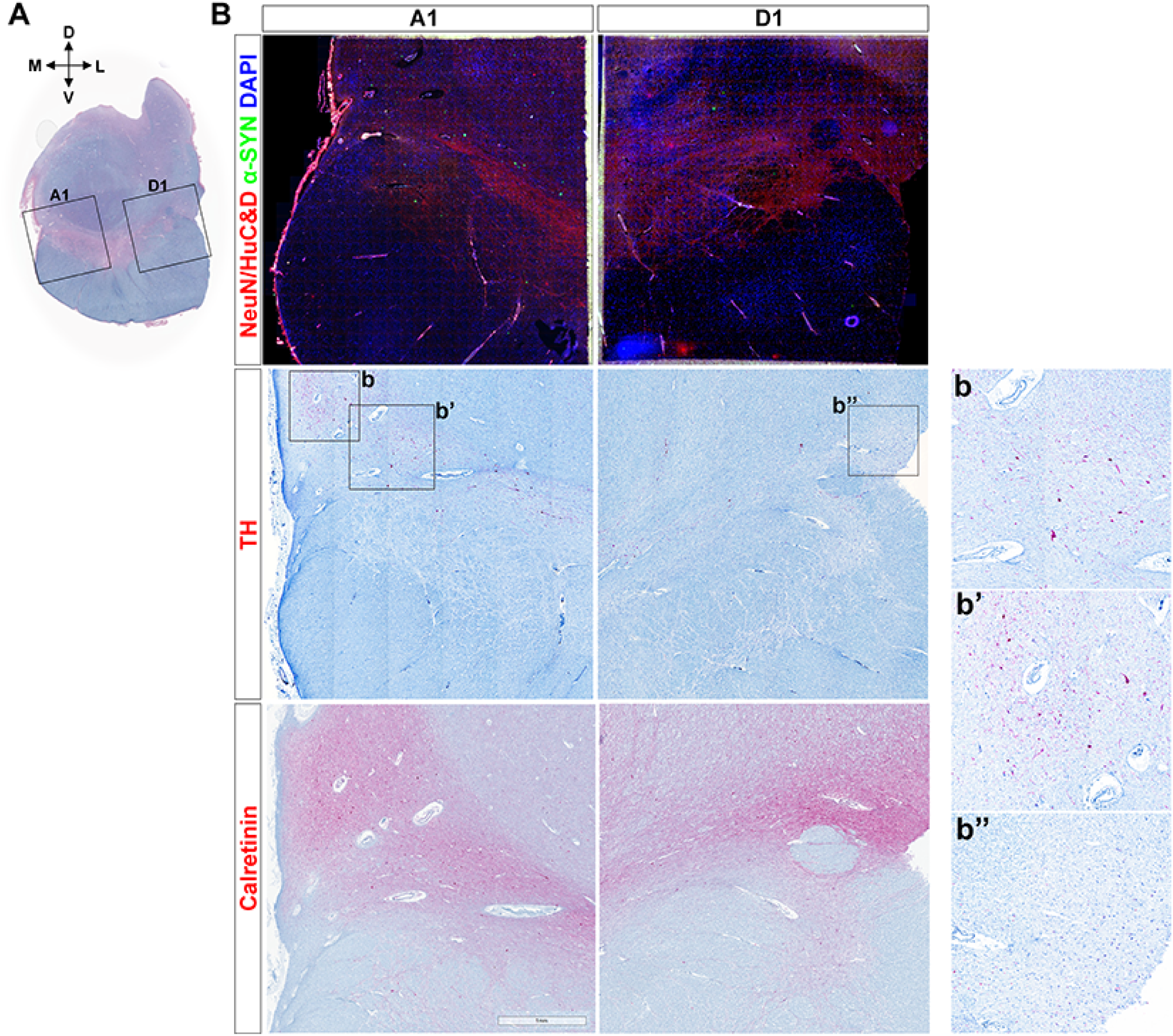
**Histologic characteristics of PD midbrain tissue used in the Visium study**. (A) A LFB staining of an adjacent section of the same left midbrain. The A1 acquisition area includes the medial portion of the SN and the D1 acquisition area includes the lateral portion of the SN, as indicated by the dotted box. (B) The immunostaining for neuronal markers, NeuN and HuC&D (red fluorescence, top panel), -SYN (green fluorescence, top panel), TH (red chromogen, middle panel) and Calretinin (red chromogen, bottom panel). The left column image of each panel shows an image corresponding to the A1 area and the right column image of each panel shows an image corresponding to the D1 area as indicated in (A). The region enriched in TH-positive dopamine neurons in A1 is enlarged in (b) and (b’), and in D1 is enlarged in (b’’).

**Supplementary Figure 2.**
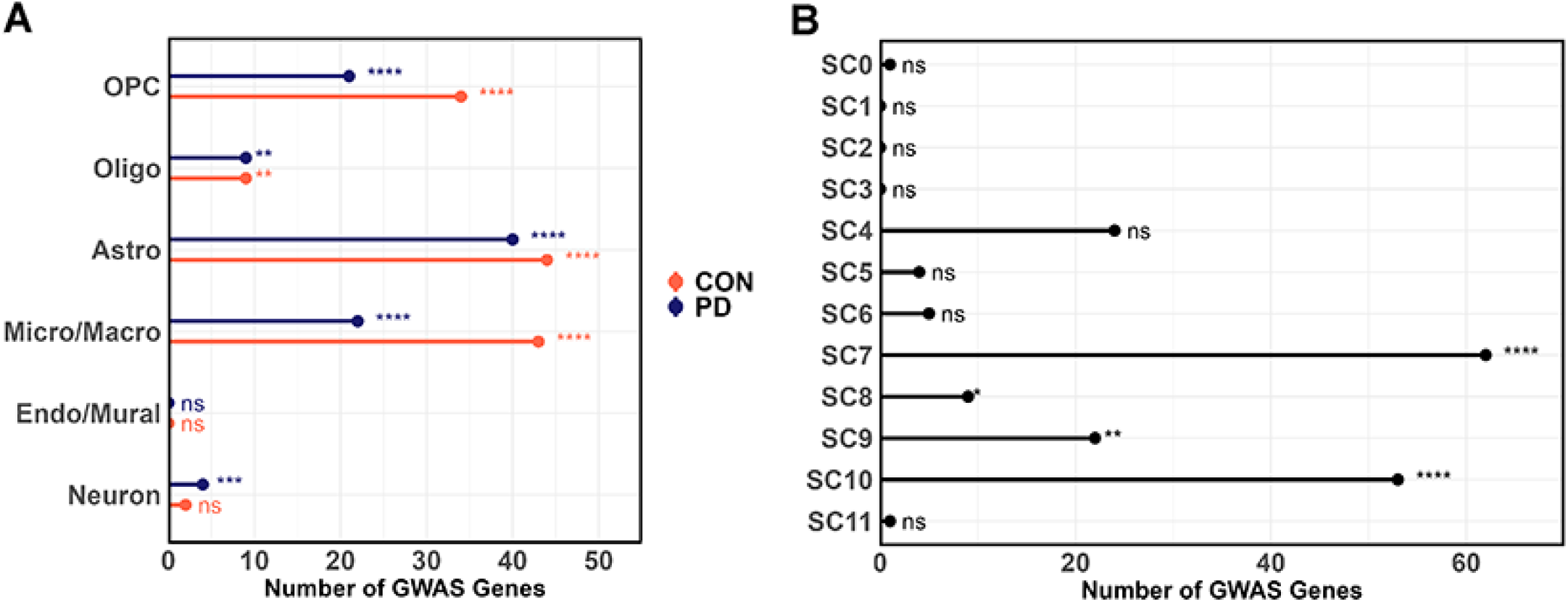
Number of GWAS genes involved in DEG per cell type and spatial cluster. (A) Count of intersecting PD GWAS genes across 12 DEG groups, differentiated between CON and PD, for 6 distinct cell types. (B) Count of intersecting PD GWAS genes across 12 spatial clusters. Asterisks depict adjusted *p*-value (corrected by *Benjamini-Hochberg* method) derived from hypergeometric test (one-sided, upper tail). ****, 0 < *p* < 0.0001; ***, 0.0001 < *p* < 0.001; **, 0.001 < *p* < 0.01; *, 0.01 < *p* < 0.05; *ns*, not significant.

**Supplementary Figure 3.**
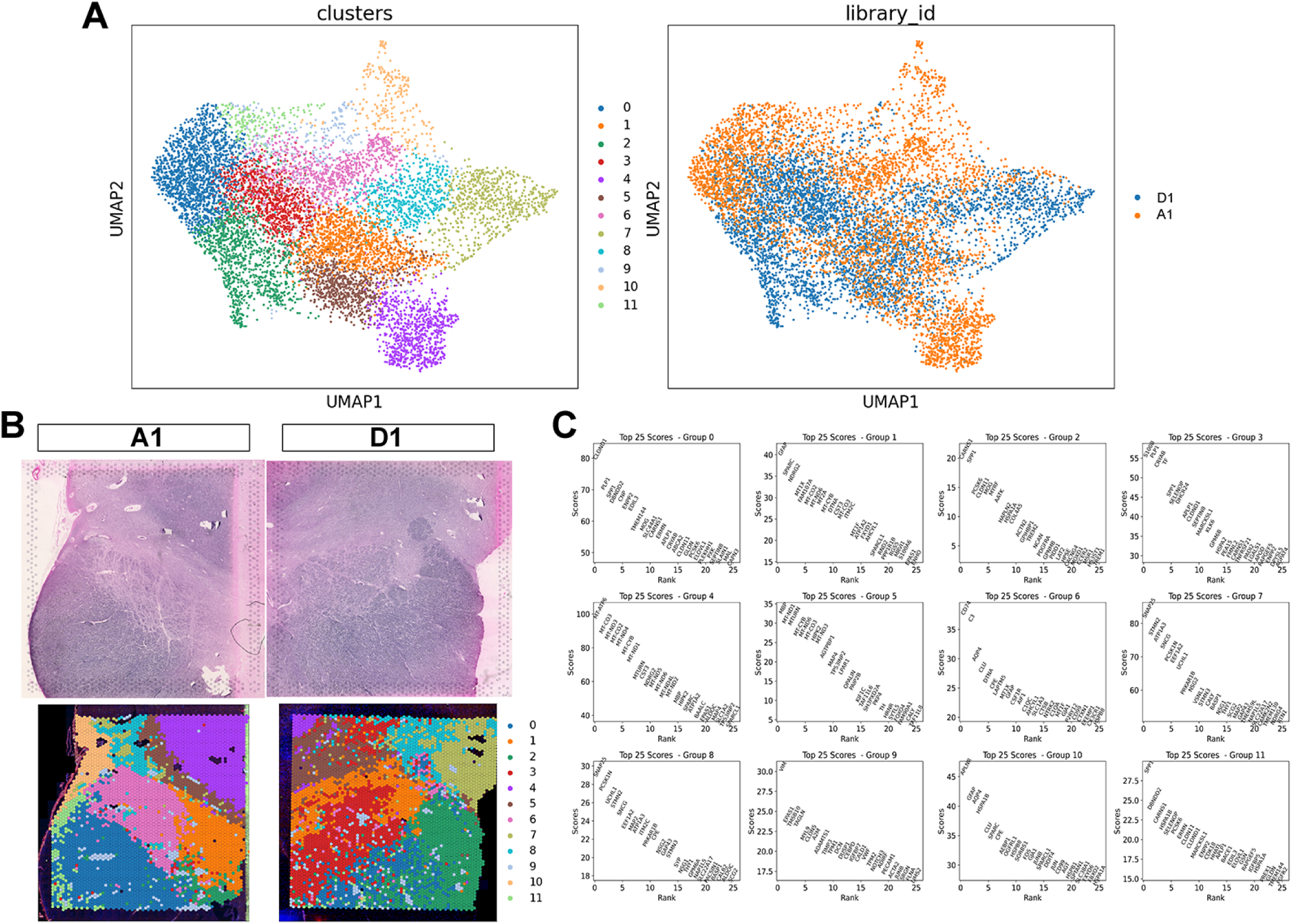
Integrated clustering between Visium spots in midbrain. (A) UMAP visualization showing twelve spatial clusters (left panel) and anatomical position of the spots (right panel). (B) Visium spots of each cluster overlaid on tissue images captured by CytAssist instruments. (C) Top 25 DEGs for each spatial cluster, ranked by score. The x-axis represents the rank, while the y-axis represents the score. Only genes with an adjusted *p*-value (corrected by the *Bonferroni-Hochberg* method) of less than 0.05 are included in the ranking.

**Supplementary Figure 4.**
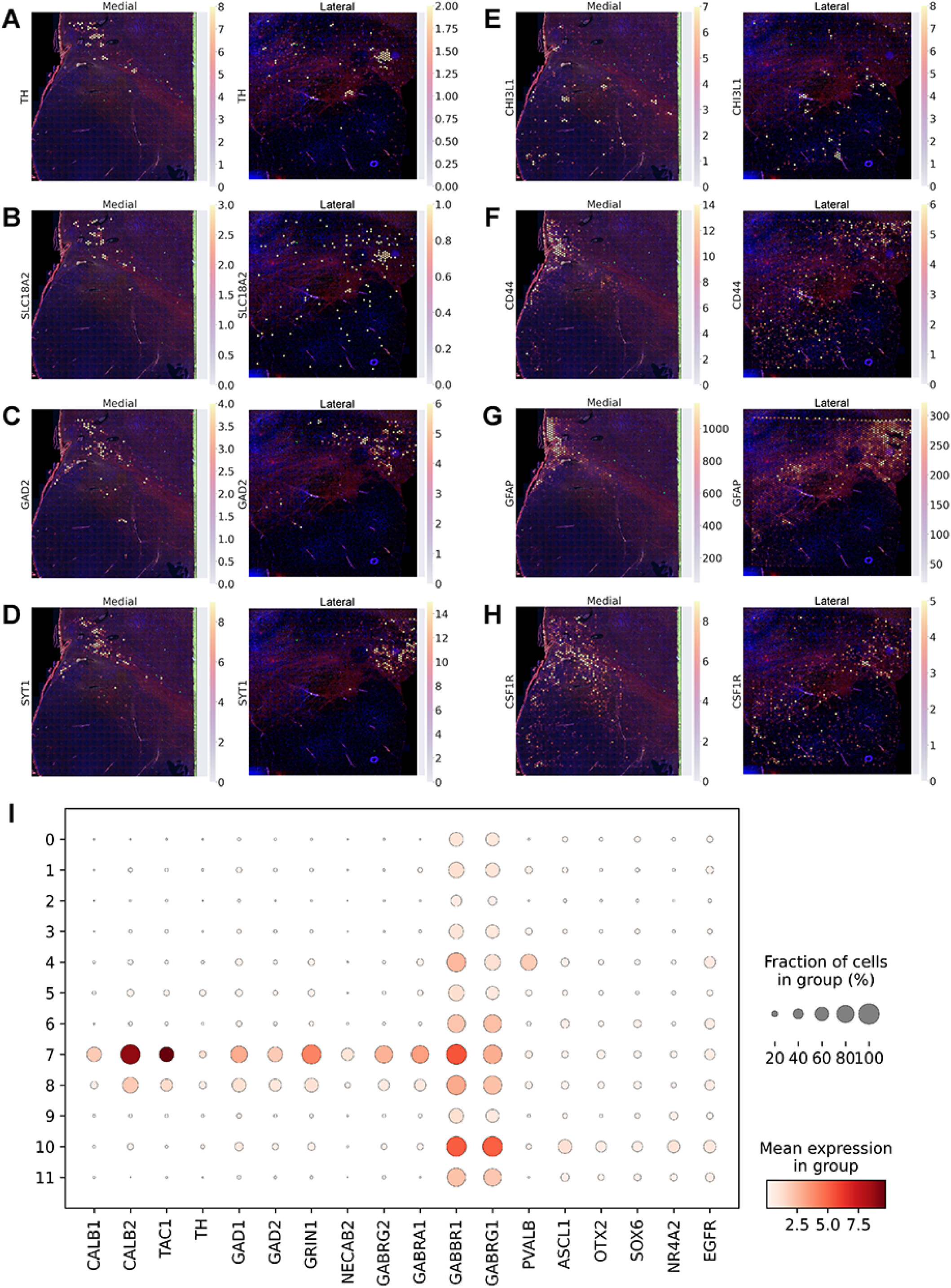
Spatial feature plots and normalized expression within each spatial cluster for genes of interest. (A - H) Normalized gene expression of spots. The left panel shows the A1 capture area (Medial), while the right panel shows the D1 capture area (Lateral). *TH* and *SLC18A2* as markers for dopaminergic neurons, *GAD2* and *SYT1* as markers for GABAergic neurons, *CHI3L1* and *CD44* as highly expressed genes in astrocytes of PD patients, *GFAP* as marker for reactive astrocytes, and *CSF1R* as marker for microglia. (I) Dot plot shows scaled gene expression for each spatial cluster. The size of the circle indicates the proportion of spots with non-zero expression.

**Supplementary Figure 5.**
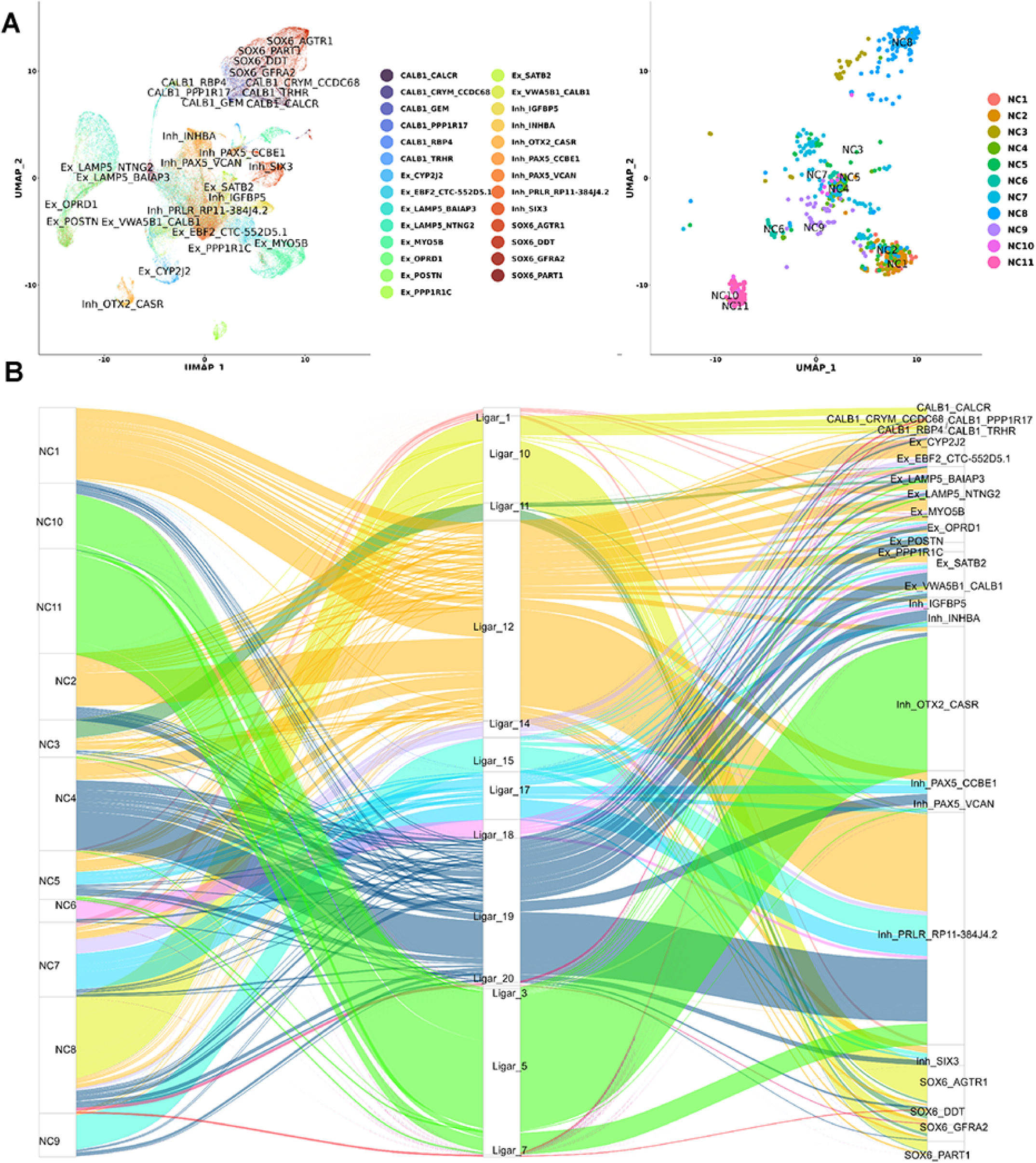
LIGER analysis between neurons from the current study data and from Kamath *et al*. data. [25]. (A) UMAP plots for public reference neurons (left panel) and the neurons obtained in this study (right panel). Both sets of data share identical UMAP coordinates. (B) Alluvial plot showing the proportional contribution of both reference data and our current data to specific LIGER clusters.

**Supplementary Figure 6.**
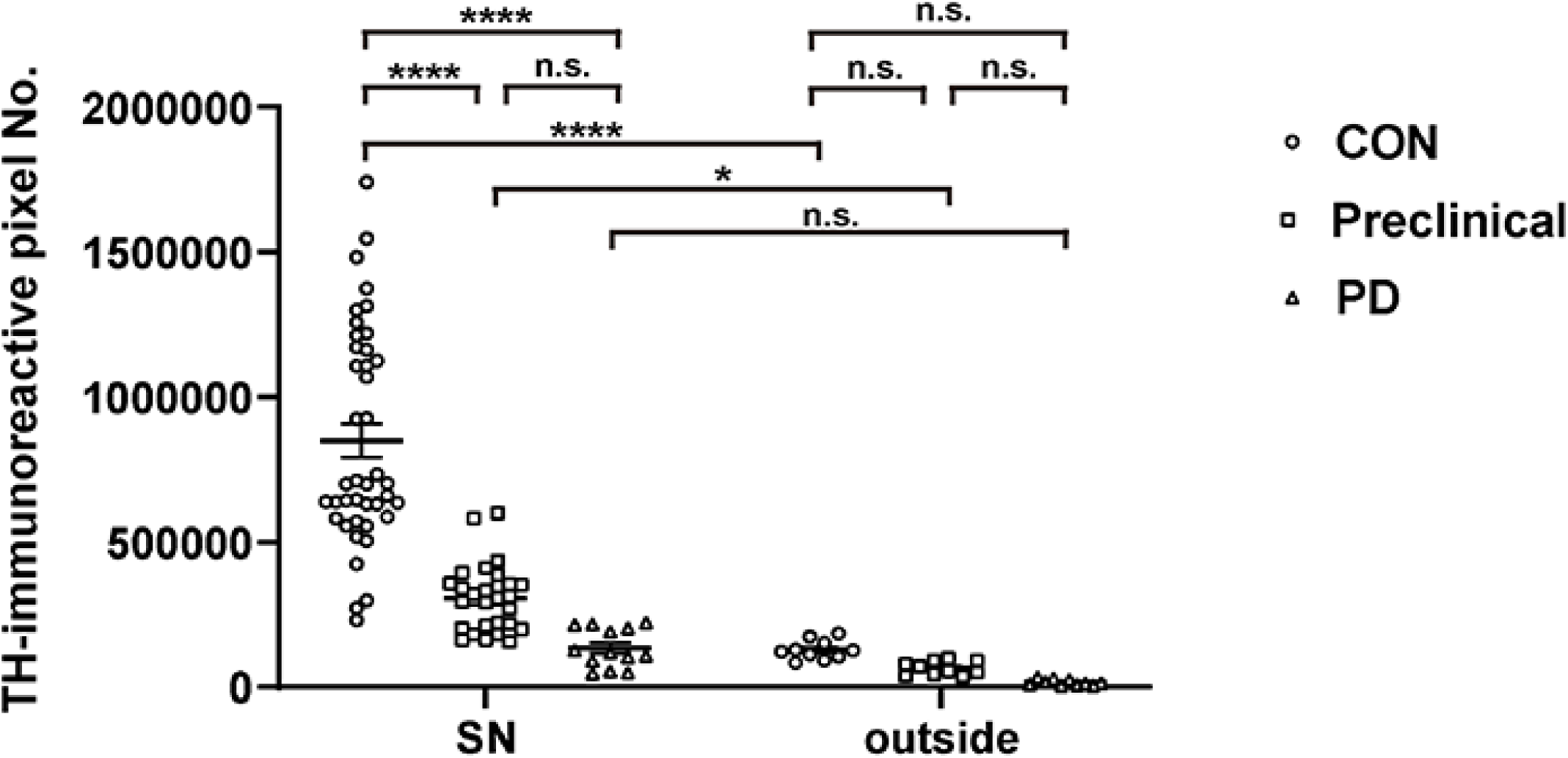
Quantification of TH-immunoreactive pixels inside the SN compared to outside. Circle point represents pixels in CON sample, square point represents pixels in preclinical PD samples, and triangle point represents pixels in PD sample. Statistics were calculated using a 2-way ANOVA followed by *Sidak*’s multiple comparison test. *, *p* < 0.05; ****, *p* < 0.0001; n.s., not significant,

**Supplementary Figure 7.**
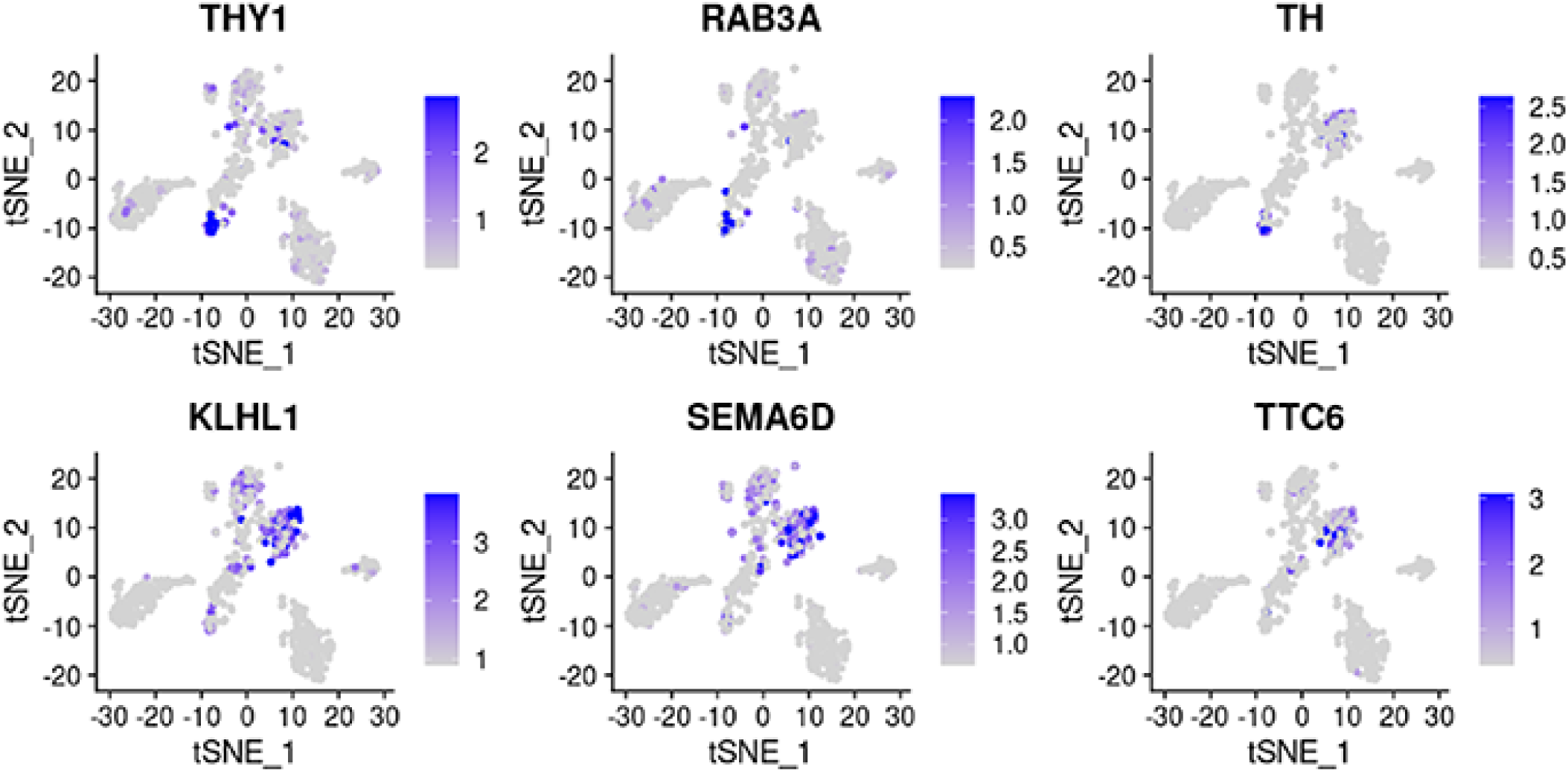
**t-SNE plot showing the normalized expression of different neuronal subtype marker genes enriched in either NC3 (upper panel) or NC8 (lower panel).**

**Supplementary Figure 8.**
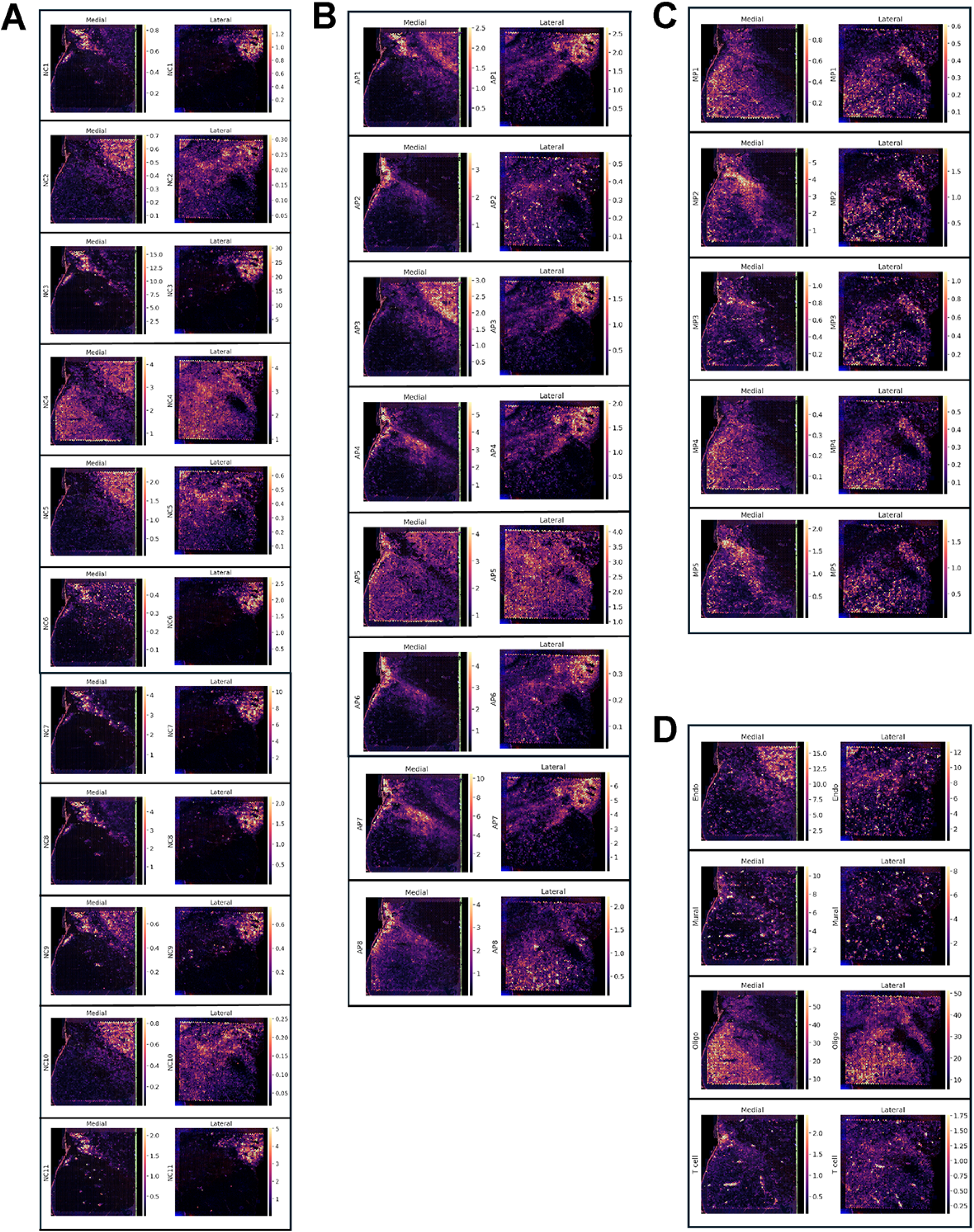
Cell2location results and visualization based on each identified subpopulation or cell type. (A) Abundance score of 11 neuronal subtypes depicted on medial (left panel) and lateral (right panel) anatomical regions. (B) Abundance score of 8 astrocyte Monocle components depicted on medial (left panel) and lateral (right panel) anatomical regions. (C) Abundance score of 5 microglia Monocle components depicted on medial (left panel) and lateral (right panel) anatomical regions. (D) Abundance score of endothelial cells, mural cells, oligodendrocytes and T-cell depicted on medial (left) and lateral (right) anatomical regions.

**Supplementary Figure 9.**
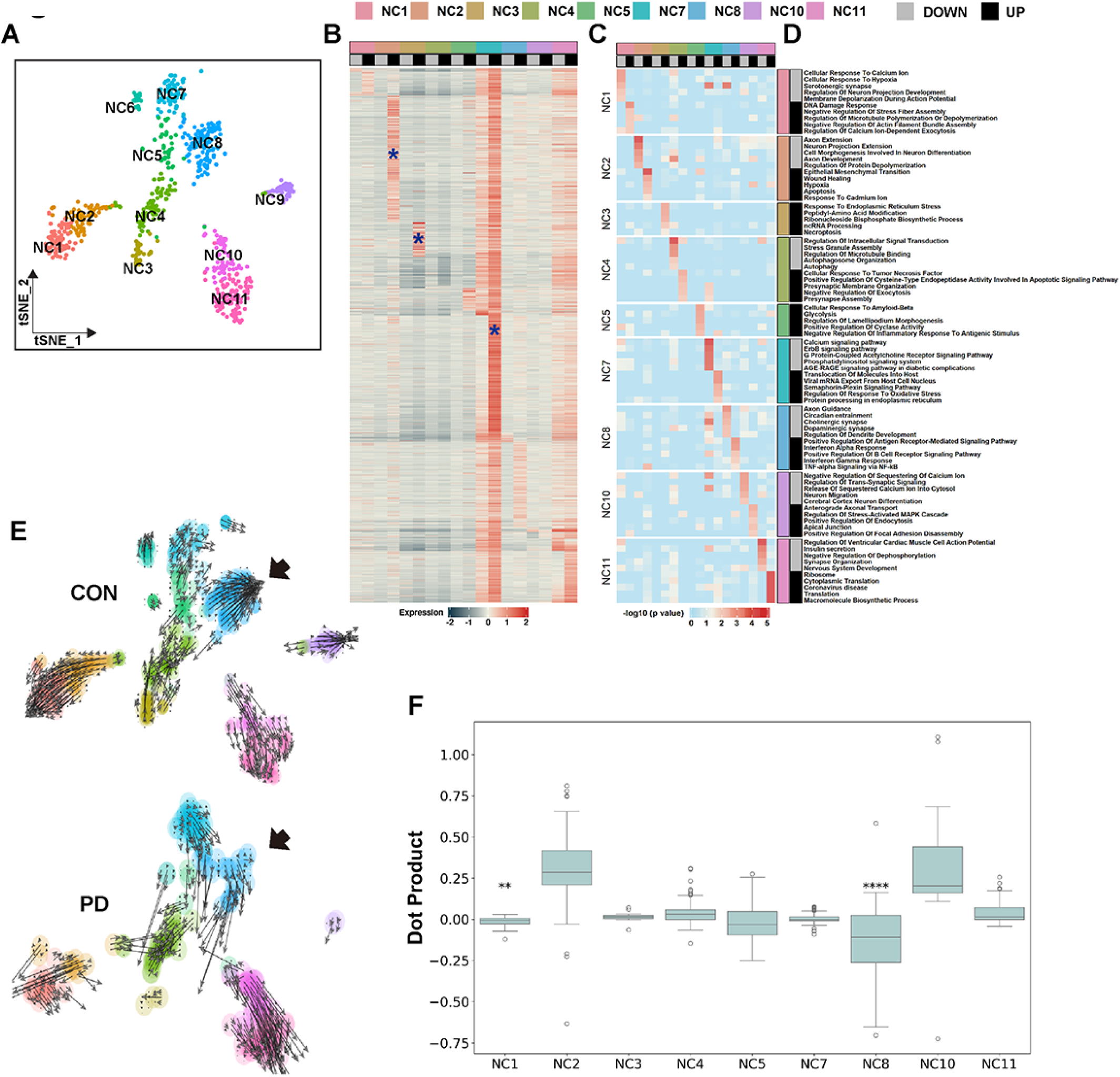
DEG and RNA velocity results in a neuronal subset. (A) tSNE plot shows 11 neuronal subtypes. (B) Heatmap of DEGs between CON and PD within each neuronal subcluster. Normalized expression is scaled for each column. (C) Heatmap of GO terms for the DEGs identified in (B). The *p*-value is represented in *-log10*. (D) Top5 GO terms corresponding to (C). (E) Embedded RNA velocity vector of both CON (upper panel) and PD (lower panel) cells. In particular, the RNA velocity vectors in NC8 are observed to be moving in the opposite direction, marked with a black arrow. (F) Boxplot showing the inner product between an embedded velocity vector of a cell and velocity vectors of nearest 3 opposite condition cells. ****, 0 < *p* < 0.0001, adjusted by *Benjamini-Hochberg* method; **, 0.01 < *p* < 0.001 (One-sided *Wilcoxon* signed rank test). Despite the fact that NC1 shows significance, the inner product of the velocity vectors across two conditions is less than 0 and its median is -0.005, which is negligible.

**Supplementary Figure 10.**
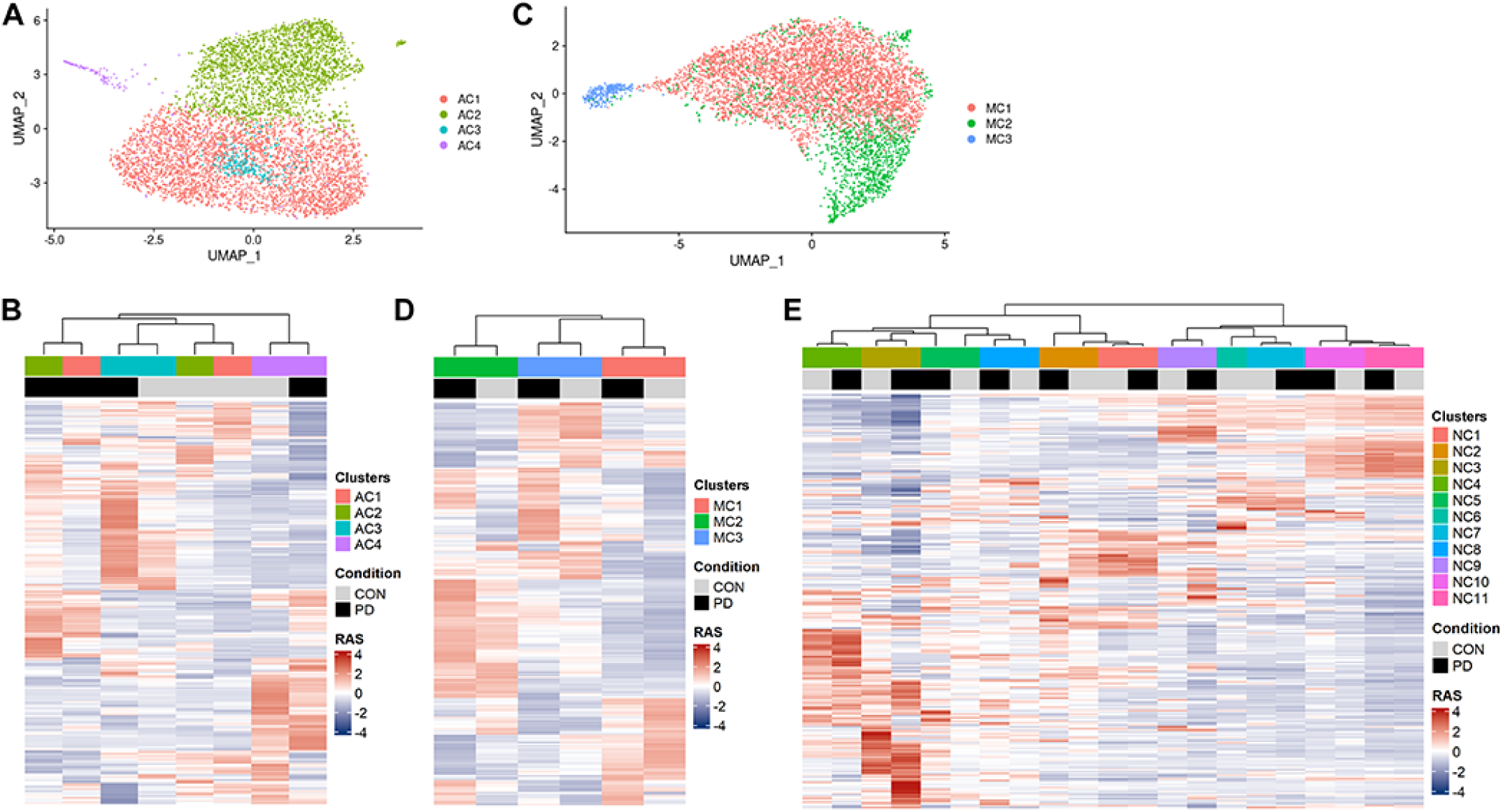
Seurat-based clustering in astrocytes and microglia, and regulon comparison between Seurat clusters for each cell type. (A, C) UMAP plots show the seurat clusters of astrocytes (A) and microglia (C), which are identified using the *Louvain* algorithm incorporated in the *Seurat* package. AC, astrocyte cluster; MC, microglia cluster. (B, D) Heatmaps of regulons over the CON and PD state of each astrocyte (B) and microglia (D) clusters. The color on the heatmap signifies the regulon activity score (RAS), which is scaled individually for each gene using SCENIC followed by hierarchical clustering. (E) Heatmap of regulons over the CON and PD state of each neuronal cluster (NC). RAS scaled for each gene, in the same way as in (D).

**Supplementary Figure 11.**
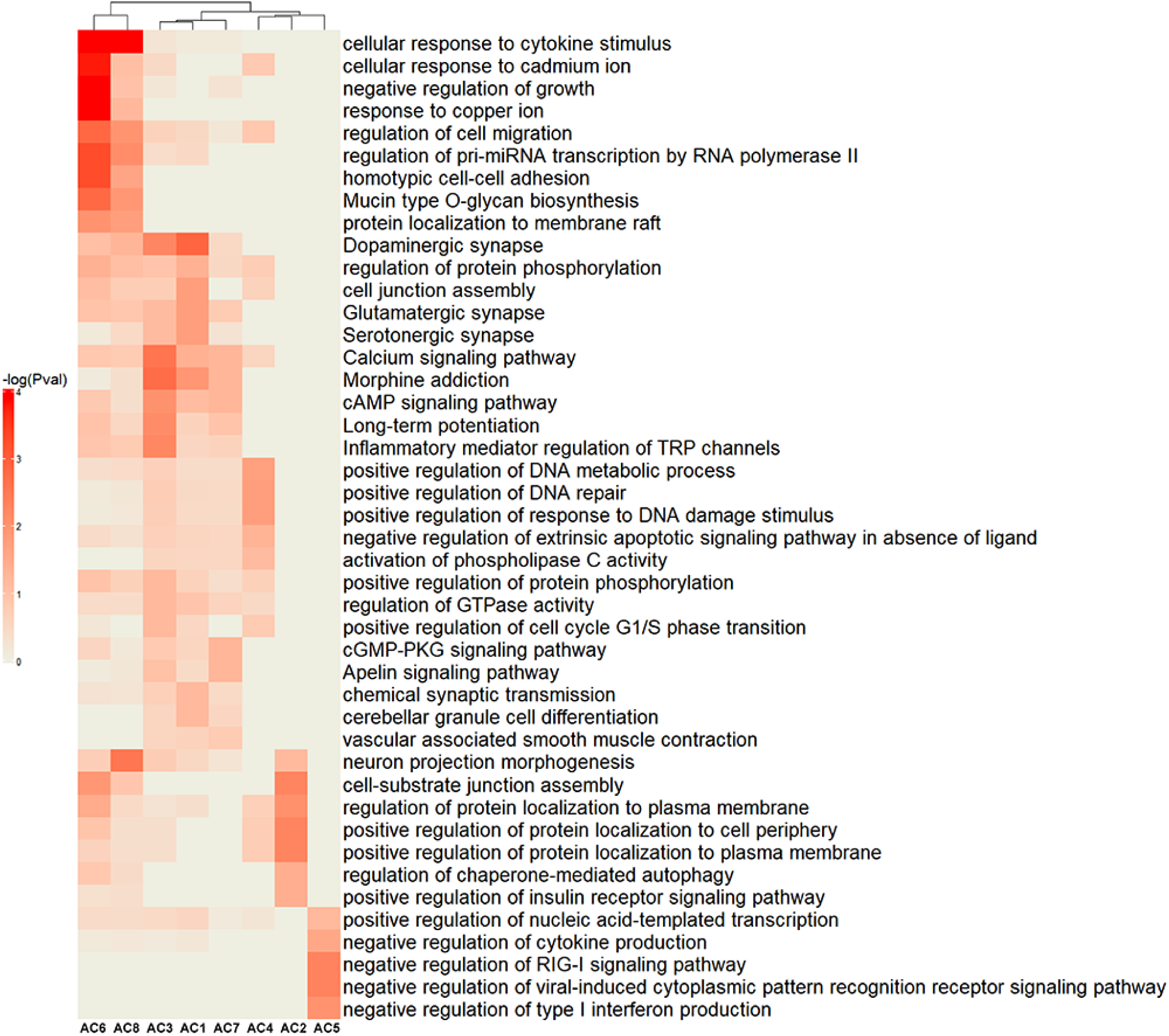
Heatmap of the enriched GO terms associated with DEGs that are upregulated in specific astrocyte components. The color on the heatmap corresponds to the *p*-value. AC, astrocyte components.

**Supplementary Figure 12.**
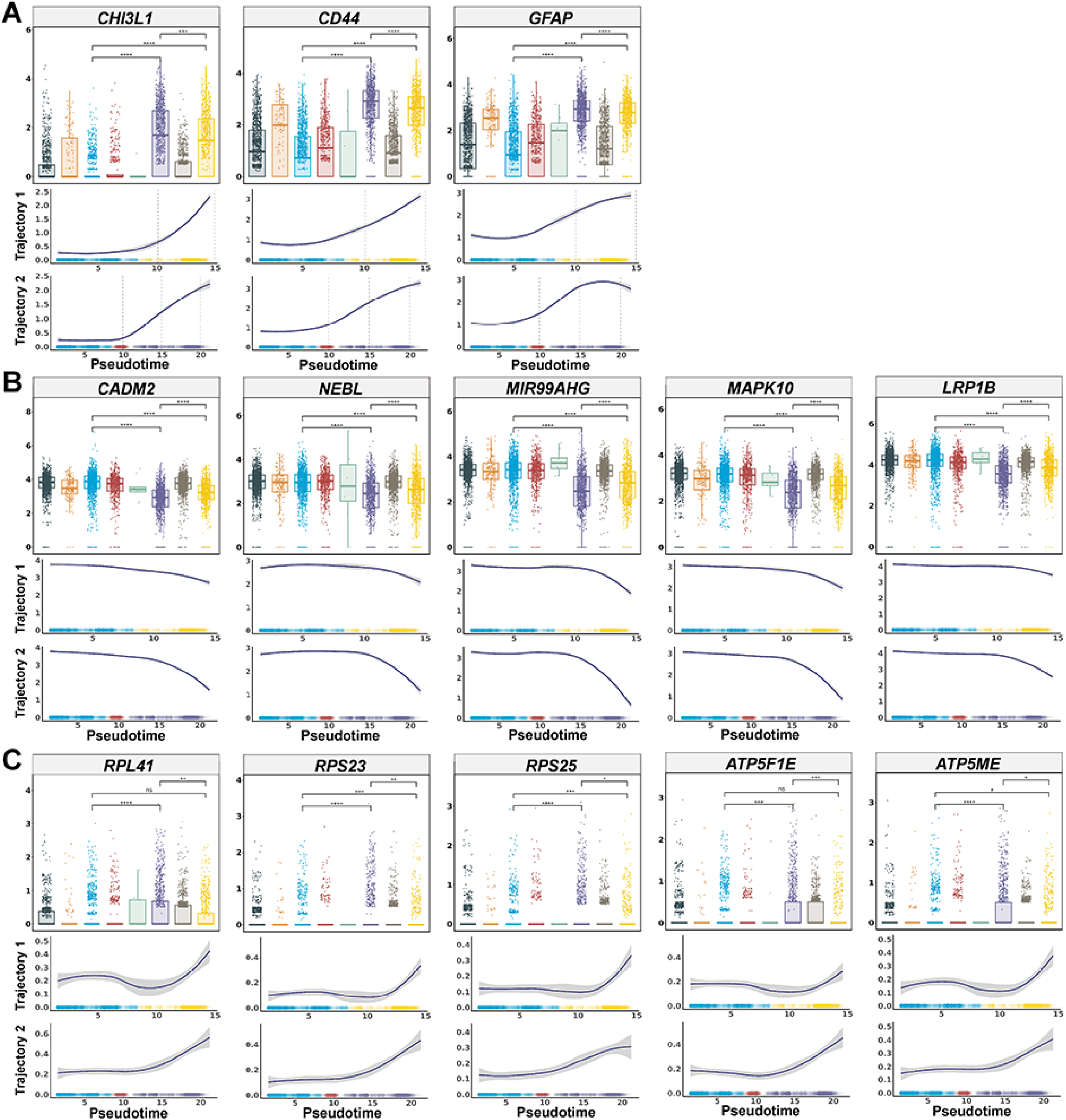
Gene expression changes between astrocyte components and over pseudotime. (A) Changes in reactive glial markers, *CHI3L1*, *CD44* and *GFAP*. (B) Changes in representative genes that show a more dramatic decrease in trajectory 2 than in trajectory 1. (C) Changes in representative genes that show a more dramatic increase in trajectory 2 than in trajectory 1. The upper panel for each gene displays a box plot of normalized gene expression grouped by astrocyte Monocle components. The colors for each component are kept consistent with the main figures such as Fig. 3B. Asterisks indicate the same information as Fig. 3I,J and K. Thus, the significance of differences, determined by the *Wilcoxon* rank-sum test, is indicated as follows: ****, 0 < *p* < 0.0001, adjusted by *Benjamini-Hochberg* method; ***, 0.0001 < *p* < 0.001; **, 0.001 < *p* < 0.01; *, 0.01 < *p* < 0.05; *ns*, not significant. The lower panel for each gene shows locally regressed expression data across pseudotime. The upper curve corresponds to Trajectory 1, while the lower curve corresponds to Trajectory 2. The gray area around the line represents the 95% confidence interval, calculated using the standard error. *ns*, not significant.

**Supplementary Figure 13.**
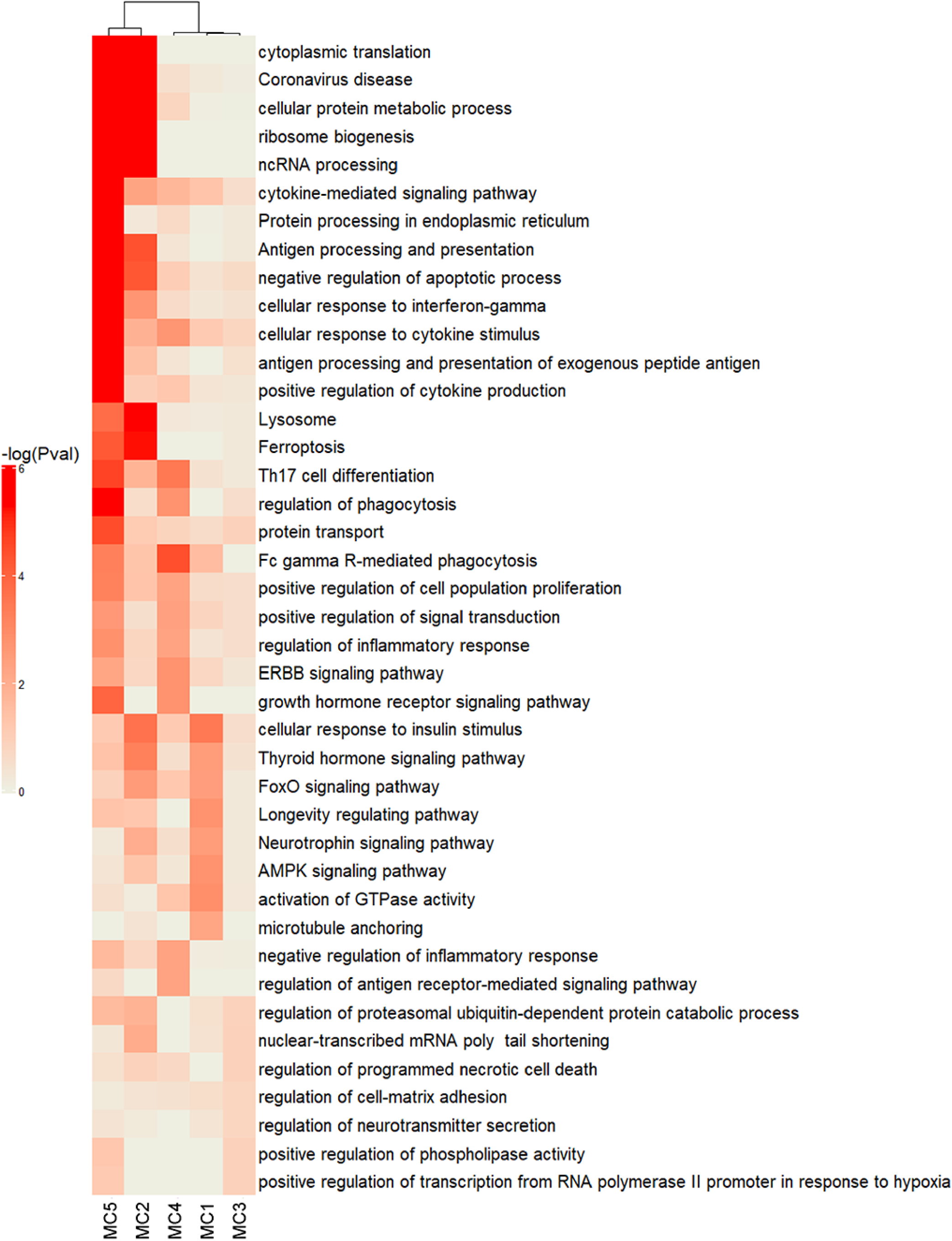
Heatmap of the enriched GO terms associated with DEGs that are upregulated in specific microglia components. The color on the heatmap corresponds to the *p*-value. MC, microglia component.

**Supplementary Figure 14.**
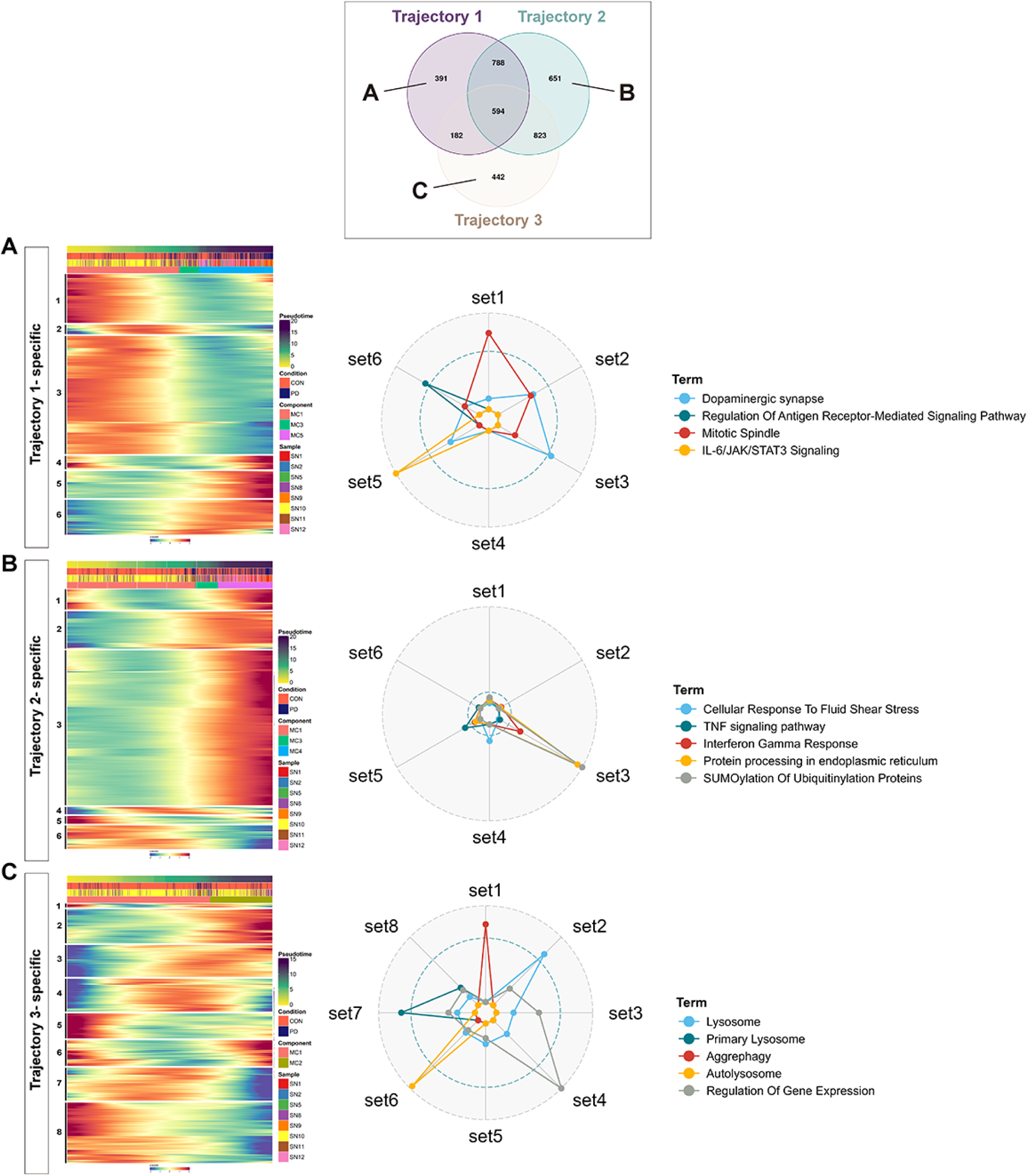
Unique monocle DEGs in individual microglia trajectories. In the Venn diagram, the number of trajectory-specific DEGs were indicated for Trajectory 1 (A), Trajectory 2 (B), and Trajectory 3 (C). Each heatmaps show the expression changes of trajectory-specific DEGs along the pseudotime. The DEGs in each gene set are shared in Supplementary Table 10_4, which are segregated into 6-8 different gene sets. Corresponding GO terms are depicted on the radar plot, representing the adjusted *p*-value for each term in relation to the gene sets, with *-log10* scaled, corrected with *Benjamini-Hochberg* method. Dotted line in the plot represents *-log10* (adjusted *p*-value). (A) Genes specific to Trajectory 1 show dominant down-regulation over pseudotime, with the largest gene set related to dopaminergic synapse and the next largest gene set associated with mitotic spindle, indicating potential PD-induced changes in microglia function or proliferation. In contrast, the upregulated Gene sets, 5 and 6, were linked to immune related terms, including regulation of antigen receptor mediated signaling pathway and IL-6/JAK/STAT3 signaling. (B) Gene specific to Trajectory 2 mainly characterized by up-regulation over pseudotime, with a significant proportion associated with SUMOylation of ubiquitinylation proteins and protein processing in ER protein modification process, as well as interferon gamma response. (C) Trajectory 3 showing serial transient changes in the expression of several Gene sets, particularly those involved in lysosome-dependent protein degradation processes, but not showing a strong association with the immune response. These distinct features provide insights into cellular changes differ in two PD-associated microglia fates and highlight the important implication of inflammatory response in PD progression.

**Supplementary Figure 15.**
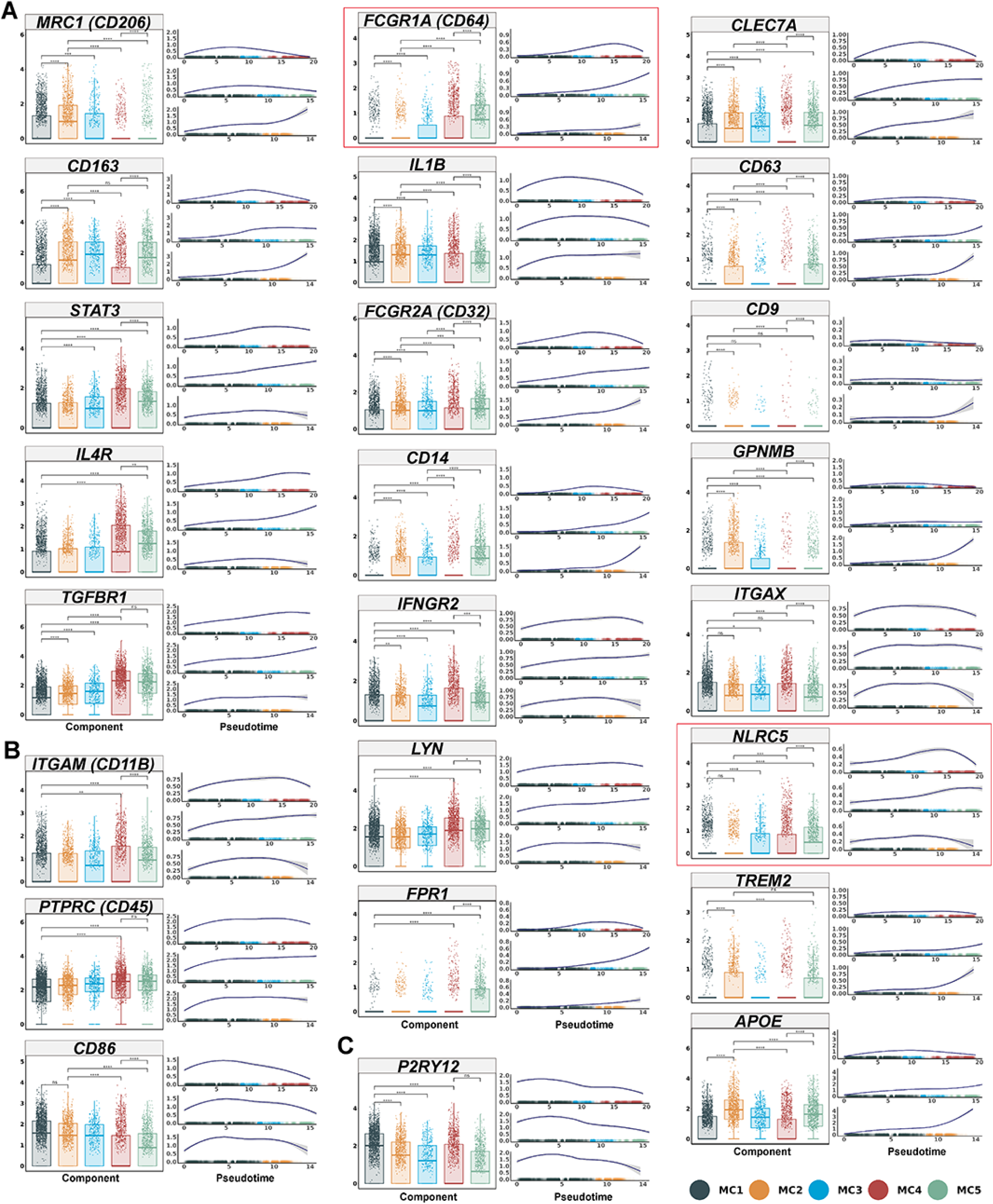
Gene expression changes between microglia components and over pseudotime. (A) Changes in known M2 microglia markers. (B) Changes in known M1 microglia markers. (C) Changes in disease-associated microglia (DAM) related genes. The left panel for each gene displays a boxplot of normalized gene expression grouped by microglia Monocle components. The right panel for each gene shows locally regressed expression data across pseudotime. The top curve is associated with Trajectory 1, the middle curve corresponds to Trajectory 2, and the bottom curve is linked to Trajectory 3. The gray area around the line represents the 95% confidence interval. The significance of differences, determined by the *Wilcoxon* rank-sum test, is indicated as follows: ****, 0 < *p* < 0.0001; ***, 0.0001 < *p* < 0.001; **, 0.001 < *p* < 0.01; *, 0.01 < *p* < 0.05. *p* indicates an adjusted *p*-value corrected by the *Benjamini-Hochberg* method. ns, not significant. The genes in the red boxes are the strong PD-associated microglial marker candidates mentioned in the discussion session.

**Supplementary Figure 16.**
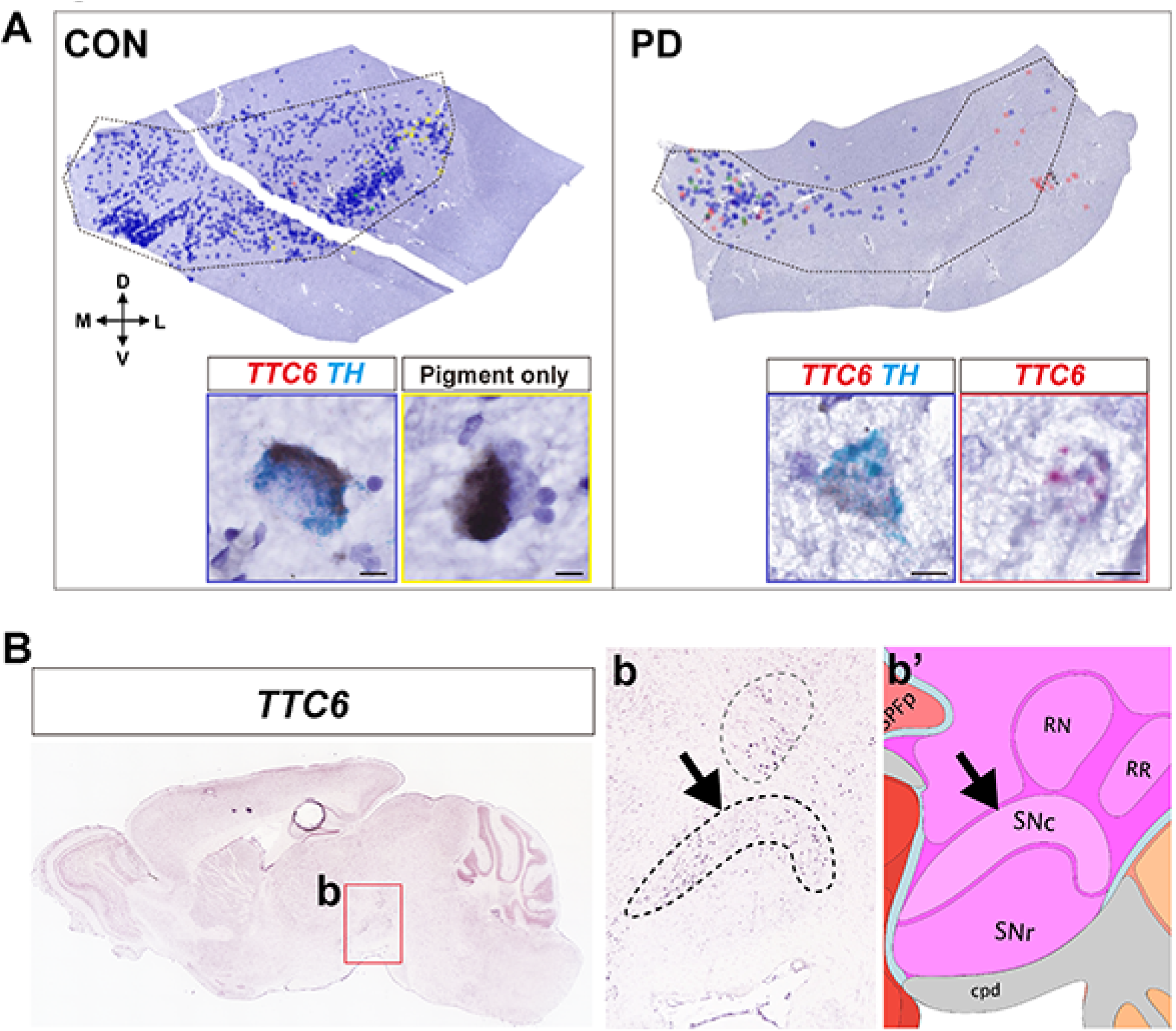
*TTC6* mRNA expression in SN neurons validated by RNAscope and verified in the Allen Brain Atlas. (A) Validation of *TCC6*-positive dopaminergic neurons in SN from CON (left panel) and PD (right panel) using RNAscope. The analyzed mRNA probe positive cells were manually marked in the enlarged SN (dotted polygon). Insets show images of representative cells for each probe. Scale bar, 50 μm. (B) *In situ* hybridization for *TTC6* expression in the Allen Brain Atlas result of postnatal day 56 male mouse brain. *TTC6* expression was detected not only in the red nucleus (RN) but also in the SN pars compacta (SNc; arrow) as shown in (b). (b’) is the reference atlas corresponding to (b). SNr, substantia nigra pars reticulata; cpd, cerebral peduncle; RR, midbrain reticular nucleus, retrorubral area.

**Supplementary Table 1**. Detailed donor information, pathology characteristics, and sample usage.

**Supplementary Table 2**. DEGs in a pooled data set. (Table 2_1) cell type-specific markers. (Table 2_2) DEGs between CON and PD per cell type. (Table 2_3) DEGs intersecting with GWAS genes.

**Supplementary Table 3**. Visium spatial cluster markers.

**Supplementary Table 4**. DEGs in neurons. (Table 4-1) Neuronal subcluster markers. (Table 4_2) DEGs between CON and PD per cluster.

**Supplementary Table 5**. Quantification of RNAscope results. Number and percentage of positive cells for each in situ probe.

**Supplementary Table 6**. T-scores of genes that have significant RNA velocity within neuronal subtypes for both CON and PD groups.

**Supplementary Table 7**. SCENIC result presented by Regulon Specificity Score (RSS) and *z*-score.

**Supplementary Table 8**. DEGs in astrocytes. (Table 8_1) Enriched genes in each astrocyte monocle component. (8-2) Monocle DEGs across each astrocyte trajectory. (8-3) DEG lists for trajectory separated by Gene sets.

**Supplementary Table 9**. Scaled RNA velocity results in astrocytes. This data corresponds to Fig. 3F, right panel. Peudotime was divided into 10 grids to study gradual changes.

**Supplementary Table 10**. DEGs in microglia. (Table 10_1) Enriched genes in each microglia monocle component. (Table 10_2) Monocle DEGs across each microglia trajectory. (Table 10_3) List of DEGs shared between trajectories, separated by Gene sets. This data corresponds to Fig. 4F. (Table 10_4) List of DEGs unique to each trajectory, separated by Gene sets. This data corresponds to sFig. 14.

**Supplementary table 11**. Detailed information of genes observed in GRN. Columns G through K show the number of targets in the source trajectory.

## Reference

1. Klein C, Westenberger A. Genetics of Parkinson’s disease. Cold Spring Harb Perspect Med. 2012;2:a008888.

2. Cherian A, Divya KP. Genetics of Parkinson’s disease. Acta Neurol Belg. 2020;120:1297–305.

3. Cho J-W, Kim S-Y, Park S-S, Jeon BS. The G2019S LRRK2 Mutation is Rare in Korean Patients with Parkinson’s Disease and Multiple System Atrophy. J Clin Neurol. 2009;5:29–32.

4. Cho J-W, Kim S-Y, Park S-S, Kim H-J, Ahn T-B, Kim J-M, et al. The G2019S LRRK2 mutation is rare in Korean patients with Parkinson’s disease. Can J Neurol Sci. 2007;34:53–5.

5. Wu Y-R, Chang K-H, Chang W-T, Hsiao Y-C, Hsu H-C, Jiang P-R, et al. Genetic variants ofLRRK2 in Taiwanese Parkinson’s disease. PLoS One. 2013;8:e82001.

6. Kalinderi K, Fidani L, Bostantjopoulou S, Katsarou Z, Kotsis A. The G2019S LRRK2 mutation is uncommon amongst Greek patients with sporadic Parkinson’s disease. Eur J Neurol. 2007;14:1088–90.

7. Monfrini E, Di Fonzo A. Leucine-Rich Repeat Kinase (LRRK2) Genetics and Parkinson’s Disease. Adv Neurobiol. 2017;14:3–30.

8. Greenwood RD. Digitalis as treatment for pulmonary comsumption, 1799. IMJ Ill Med J. 1975;148:531.

9. Kingwell K. LRRK2-targeted Parkinson disease drug advances into phase III. Nat Rev Drug Discov. 2023;22:3–5.

10. Kingwell K. LRRK2 inhibitor progresses for Parkinson disease. Nat. Rev. Drug Discov. 2022. p. 558.

11. Cheon S-M, Chan L, Chan DKY, Kim JW. Genetics of Parkinson’s disease - a clinical perspective. J Mov Disord. 2012;5:33–41.

12. Ben-Joseph A, Marshall CR, Lees AJ, Noyce AJ. Ethnic Variation in the Manifestation of Parkinson’s Disease: A Narrative Review. J Parkinsons Dis. 2020;10:31–45.

13. Kurtzke JF, Goldberg ID. Parkinsonism death rates by race, sex, and geography. Neurology. 1988;38:1558–61.

14. Foo JN, Chew EGY, Chung SJ, Peng R, Blauwendraat C, Nalls MA, et al. Identification of Risk Loci for Parkinson Disease in Asians and Comparison of Risk Between Asians and Europeans: A Genome-Wide Association Study. JAMA Neurol. 2020;77:746–54.

15. Pan H, Liu Z, Ma J, Li Y, Zhao Y, Zhou X, et al. Genome-wide association study using whole-genome sequencing identifies risk loci for Parkinson’s disease in Chinese population. NPJ Parkinsons Dis. 2023;9:22.

16. Foo JN, Tan LC, Irwan ID, Au W-L, Low HQ, Prakash K-M, et al. Genome-wide association study of Parkinson’s disease in East Asians. Hum Mol Genet. 2017;26:226–32.

17. Kim JJ, Vitale D, Otani DV, Lian MM, Heilbron K, 23andMe Research Team, et al. Multi-ancestry genome-wide association meta-analysis of Parkinson’s disease. Nat Genet. 2024;56:27–36.

18. Lim S-Y, Tan AH, Ahmad-Annuar A, Klein C, Tan LCS, Rosales RL, et al. Parkinson’s disease in the Western Pacific Region. Lancet Neurol. 2019;18:865–79.

19. Chung SJ, Jung Y, Hong M, Kim MJ, You S, Kim YJ, et al. Alzheimer’s disease and Parkinson’s disease genome-wide association study top hits and risk of Parkinson’s disease in Korean population. Neurobiol Aging. 2013;34:2695.e1–7.

20. Park KW, Ryu H-S, Shin E, Park Y, Jeon SR, Kim SY, et al. Ethnicity- and sex-specific genome wide association study on Parkinson’s disease. NPJ Parkinsons Dis. 2023;9:141.

21. Agarwal D, Sandor C, Volpato V, Caffrey TM, Monzón-Sandoval J, Bowden R, et al. A single-cell atlas of the human substantia nigra reveals cell-specific pathways associated with neurological disorders. Nat Commun. 2020;11:4183.

22. Wang Q, Wang M, Choi I, Sarrafha L, Liang M, Ho L, et al. Molecular profiling of human substantia nigra identifies diverse neuron types associated with vulnerability in Parkinson’s disease. Sci Adv. 2024;10:eadi8287.

23. Smajić S, Prada-Medina CA, Landoulsi Z, Ghelfi J, Delcambre S, Dietrich C, et al. Single-cell sequencing of human midbrain reveals glial activation and a Parkinson-specific neuronal state. Brain. 2022;145:964–78.

24. Martirosyan A, Ansari R, Pestana F, Hebestreit K, Gasparyan H, Aleksanyan R, et al. Unravelling cell type-specific responses to Parkinson’s Disease at single cell resolution. Mol Neurodegener. 2024;19:7.

25. Kamath T, Abdulraouf A, Burris SJ, Langlieb J, Gazestani V, Nadaf NM, et al. Single-cell genomic profiling of human dopamine neurons identifies a population that selectively degenerates in Parkinson’s disease. Nat Neurosci. 2022;25:588–95.

26. Lee AJ, Kim C, Park S, Joo J, Choi B, Yang D, et al. Characterization of altered molecular mechanisms in Parkinson’s disease through cell type-resolved multiomics analyses. Sci Adv. 2023;9:eabo2467.

27. Fernandes HJR, Patikas N, Foskolou S, Field SF, Park J-E, Byrne ML, et al. Single-Cell Transcriptomics of Parkinson’s Disease Human In Vitro Models Reveals Dopamine Neuron-Specific Stress Responses. Cell Rep. 2020;33:108263.

28. Crary JF, Trojanowski JQ, Schneider JA, Abisambra JF, Abner EL, Alafuzoff I, et al. Primary age-related tauopathy (PART): a common pathology associated with human aging. Acta Neuropathol. 2014;128:755–66.

29. McMillan CT, Lee EB, Jefferson-George K, Naj A, Van Deerlin VM, Trojanowski JQ, et al. Alzheimer’s genetic risk is reduced in primary age-related tauopathy: a potential model of resistance? Ann Clin Transl Neurol. 2018;5:927–34.

30. Shim Y-M, Kim S-I, Lim SD, Lee K, Kim EE, Won JK, et al. An Autopsy-proven Case-based Review of Autoimmune Encephalitis. Exp Neurobiol. 2024;33:1–17.

31. Parkkinen L, Pirttilä T, Alafuzoff I. Applicability of current staging/categorization of alpha-synuclein pathology and their clinical relevance. Acta Neuropathol. 2008;115:399–407.

32. Braak H, Alafuzoff I, Arzberger T, Kretzschmar H, Del Tredici K. Staging of Alzheimer disease-associated neurofibrillary pathology using paraffin sections and immunocytochemistry. Acta Neuropathol. 2006;112:389–404.

33. Hao Y, Hao S, Andersen-Nissen E, Mauck WM 3rd, Zheng S, Butler A, et al. Integrated analysis of multimodal single-cell data. Cell. 2021;184:3573–87.e29.

34. Wang S, Sun S. Translation dysregulation in neurodegenerative diseases: a focus on ALS. Mol Neurodegener. 2023;18:58.

35. Vanni S, Zattoni M, Moda F, Giaccone G, Tagliavini F, Haïk S, et al. Hemoglobin mRNA Changes in the Frontal Cortex of Patients with Neurodegenerative Diseases. Front Neurosci. 2018;12:8.

36. Ferrer-Raventós P, Beyer K. Alternative platelet activation pathways and their role in neurodegenerative diseases. Neurobiol Dis. 2021;159:105512.

37. McGinnis CS, Murrow LM, Gartner ZJ. DoubletFinder: Doublet Detection in Single-Cell RNA Sequencing Data Using Artificial Nearest Neighbors. Cell Syst. 2019;8:329–37.e4.

38. Stuart T, Butler A, Hoffman P, Hafemeister C, Papalexi E, Mauck WM 3rd, et al. Comprehensive Integration of Single-Cell Data. Cell. 2019;177:1888–902.e21.

39. Welch JD, Kozareva V, Ferreira A, Vanderburg C, Martin C, Macosko EZ. Single-Cell Multi-omic Integration Compares and Contrasts Features of Brain Cell Identity. Cell. 2019;177:1873–87.e17.

40. Simón-Sánchez J, Schulte C, Bras JM, Sharma M, Gibbs JR, Berg D, et al. Genome-wide association study reveals genetic risk underlying Parkinson’s disease. Nat Genet. 2009;41:1308–12.

41. Nalls MA, Blauwendraat C, Vallerga CL, Heilbron K, Bandres-Ciga S, Chang D, et al. Identification of novel risk loci, causal insights, and heritable risk for Parkinson’s disease: a meta-analysis of genome-wide association studies. Lancet Neurol. 2019;18:1091–102.

42. Iwaki H, Blauwendraat C, Leonard HL, Kim JJ, Liu G, Maple-Grødem J, et al. Genomewide association study of Parkinson’s disease clinical biomarkers in 12 longitudinal patients’ cohorts. Mov Disord. 2019;34:1839–50.

43. Gu X-J, Su W-M, Dou M, Jiang Z, Duan Q-Q, Yin K-F, et al. Expanding causal genes for Parkinson’s disease via multi-omics analysis. NPJ Parkinsons Dis. 2023;9:146.

44. Chang D, Nalls MA, Hallgrímsdóttir IB, Hunkapiller J, van der Brug M, Cai F, et al. A meta-analysis of genome-wide association studies identifies 17 new Parkinson’s disease risk loci. Nat Genet. 2017;49:1511–6.

45. Makarious MB, Lake J, Pitz V, Ye Fu A, Guidubaldi JL, Solsberg CW, et al. Large-scale rare variant burden testing in Parkinson’s disease. Brain. 2023;146:4622–32.

46. Li B, Zhao G, Zhou Q, Xie Y, Wang Z, Fang Z, et al. Gene4PD: A Comprehensive Genetic Database of Parkinson’s Disease. Front Neurosci. 2021;15:679568.

47. Hie B, Bryson B, Berger B. Efficient integration of heterogeneous single-cell transcriptomes using Scanorama. Nat Biotechnol. 2019;37:685–91.

48. Kleshchevnikov V, Shmatko A, Dann E, Aivazidis A, King HW, Li T, et al. Cell2location maps fine-grained cell types in spatial transcriptomics. Nat Biotechnol. 2022;40:661–71.

49. Qiu X, Hill A, Packer J, Lin D, Ma Y-A, Trapnell C. Single-cell mRNA quantification and differential analysis with Census. Nat Methods. 2017;14:309–15.

50. Bergen V, Lange M, Peidli S, Wolf FA, Theis FJ. Generalizing RNA velocity to transient cell states through dynamical modeling. Nat Biotechnol. 2020;38:1408–14.

51. Xie Z, Bailey A, Kuleshov MV, Clarke DJB, Evangelista JE, Jenkins SL, et al. Gene Set Knowledge Discovery with Enrichr. Curr Protoc. 2021;1:e90.

52. Aibar S, González-Blas CB, Moerman T, Huynh-Thu VA, Imrichova H, Hulselmans G, et al. SCENIC: single-cell regulatory network inference and clustering. Nat Methods. 2017;14:1083–6.

53. Kim J, T Jakobsen S, Natarajan KN, Won K-J. TENET: gene network reconstruction using transfer entropy reveals key regulatory factors from single cell transcriptomic data. Nucleic Acids Res. 2021;49:e1.

54. Shannon P, Markiel A, Ozier O, Baliga NS, Wang JT, Ramage D, et al. Cytoscape: a software environment for integrated models of biomolecular interaction networks. Genome Res. 2003;13:2498–504.

55. Morris JH, Apeltsin L, Newman AM, Baumbach J, Wittkop T, Su G, et al. clusterMaker: a multi-algorithm clustering plugin for Cytoscape. BMC Bioinformatics. 2011;12:436.

56. Bankhead P, Loughrey MB, Fernández JA, Dombrowski Y, McArt DG, Dunne PD, et al. QuPath: Open source software for digital pathology image analysis. Sci Rep. 2017;7:16878.

57. Van Hove H, Martens L, Scheyltjens I, De Vlaminck K, Pombo Antunes AR, De Prijck S, et al. A single-cell atlas of mouse brain macrophages reveals unique transcriptional identities shaped by ontogeny and tissue environment. Nat Neurosci. 2019;22:1021–35.

58. Masuda T, Amann L, Prinz M. Novel insights into the origin and development of CNS macrophage subsets. Clin Transl Med. 2022;12:e1096.

59. Tredicine M, Camponeschi C, Pirolli D, Lucchini M, Valentini M, Geloso MC, et al. A TLR/CD44 axis regulates T cell trafficking in experimental and human multiple sclerosis. iScience. 2022;25:103763.

60. Korin B, Ben-Shaanan TL, Schiller M, Dubovik T, Azulay-Debby H, Boshnak NT, et al. High-dimensional, single-cell characterization of the brain’s immune compartment. Nat Neurosci. 2017;20:1300–9.

61. Chiba H, Ichikawa-Tomikawa N, Imura T, Sugimoto K. The region-selective regulation of endothelial claudin-5 expression and signaling in brain health and disorders. J Cell Physiol. 2021;236:7134–43.

62. Bondjers C, He L, Takemoto M, Norlin J, Asker N, Hellström M, et al. Microarray analysis of blood microvessels from PDGF-B and PDGF-Rbeta mutant mice identifies novel markers for brain pericytes. FASEB J. 2006;20:1703–5.

63. Lin J-Z, Duan M-R, Lin N, Zhao W-J. The emerging role of the chondroitin sulfate proteoglycan family in neurodegenerative diseases. Rev Neurosci. 2021;32:737–50.

64. Pintér P, Alpár A. The Role of Extracellular Matrix in Human Neurodegenerative Diseases. Int J Mol Sci [Internet]. 2022;23. Available from: 10.3390/ijms231911085

65. Burgess RW, Crish SD. Editorial: Axonopathy in Neurodegenerative Disease. Front Neurosci. 2018;12:769.

66. Lee CR, Tepper JM. Morphological and physiological properties of parvalbumin- and calretinin-containing gamma-aminobutyric acidergic neurons in the substantia nigra. J Comp Neurol. 2007;500:958–72.

67. Barraviera B, Mendes RP, Pereira PC, Machado JM, Curi PR, Meira DA. Measurement of glucose-6-phosphate dehydrogenase and glutathione reductase activity in patients with paracoccidioidomycosis treated with ketoconazole. Mycopathologia. 1988;104:87–91.

68. Tepper JM, Martin LP, Anderson DR. GABAA receptor-mediated inhibition of rat substantia nigra dopaminergic neurons by pars reticulata projection neurons. J Neurosci. 1995;15:3092–103.

69. Alharbi B, Al-Kuraishy HM, Al-Gareeb AI, Elekhnawy E, Alharbi H, Alexiou A, et al. Role of GABA pathway in motor and non-motor symptoms in Parkinson’s disease: a bidirectional circuit. Eur J Med Res. 2024;29:205.

70. Błaszczyk JW. Parkinson’s Disease and Neurodegeneration: GABA-Collapse Hypothesis. Front Neurosci. 2016;10:269.

71. Jonas A, Thiem S, Kuhlmann T, Wagener R, Aszodi A, Nowell C, et al. Axonally derived matrilin-2 induces proinflammatory responses that exacerbate autoimmune neuroinflammation. J Clin Invest. 2014;124:5042–56.

72. Esteves AR, Arduíno DM, Swerdlow RH, Oliveira CR, Cardoso SM. Microtubule depolymerization potentiates alpha-synuclein oligomerization. Front Aging Neurosci. 2010;1:5.

73. Kim S, Kim DK, Jeong S, Lee J. The Common Cellular Events in the Neurodegenerative Diseases and the Associated Role of Endoplasmic Reticulum Stress. Int J Mol Sci [Internet]. 2022;23. Available from: 10.3390/ijms23115894

74. Matus S, Glimcher LH, Hetz C. Protein folding stress in neurodegenerative diseases: a glimpse into the ER. Curr Opin Cell Biol. 2011;23:239–52.

75. Sun Y, Vashisht AA, Tchieu J, Wohlschlegel JA, Dreier L. Voltage-dependent anion channels (VDACs) recruit Parkin to defective mitochondria to promote mitochondrial autophagy. J Biol Chem. 2012;287:40652–60.

76. Morishita H, Eguchi T, Tsukamoto S, Sakamaki Y, Takahashi S, Saito C, et al. Organelle degradation in the lens by PLAAT phospholipases. Nature. 2021;592:634–8.

77. Staring J, von Castelmur E, Blomen VA, van den Hengel LG, Brockmann M, Baggen J, et al. PLA2G16 represents a switch between entry and clearance of Picornaviridae. Nature. 2017;541:412–6.

78. Lovero KL, Fukata Y, Granger AJ, Fukata M, Nicoll RA. The LGI1-ADAM22 protein complex directs synapse maturation through regulation of PSD-95 function. Proc Natl Acad Sci U S A. 2015;112:E4129– 37.

79. Zhang Z-H, Song G-L. Roles of Selenoproteins in Brain Function and the Potential Mechanism of Selenium in Alzheimer’s Disease. Front Neurosci. 2021;15:646518.

80. Bellinger FP, Raman AV, Rueli RH, Bellinger MT, Dewing AS, Seale LA, et al. Changes in selenoprotein P in substantia nigra and putamen in Parkinson’s disease. J Parkinsons Dis. 2012;2:115– 26.

81. Kim H, Wu X, Lee J. SLC31 (CTR) family of copper transporters in health and disease. Mol Aspects Med. 2013;34:561–70.

82. Birger A, Ottolenghi M, Perez L, Reubinoff B, Behar O. ALS-related human cortical and motor neurons survival is differentially affected by Sema3A. Cell Death Dis. 2018;9:256.

83. Good PF, Alapat D, Hsu A, Chu C, Perl D, Wen X, et al. A role for semaphorin 3A signaling in the degeneration of hippocampal neurons during Alzheimer’s disease. J Neurochem. 2004;91:716–36.

84. Bifsha P, Yang J, Fisher RA, Drouin J. Rgs6 is required for adult maintenance of dopaminergic neurons in the ventral substantia nigra. PLoS Genet. 2014;10:e1004863.

85. Lo P-S, Rymar VV, Kennedy TE, Sadikot AF. The netrin-1 receptor DCC promotes the survival of a subpopulation of midbrain dopaminergic neurons: Relevance for ageing and Parkinson’s disease. J Neurochem. 2022;161:254–65.

86. Jasmin M, Ahn EH, Voutilainen MH, Fombonne J, Guix C, Viljakainen T, et al. Netrin-1 and its receptor DCC modulate survival and death of dopamine neurons and Parkinson’s disease features. EMBO J. 2021;40:e105537.

87. Hegarty SV, Sullivan AM, O’Keeffe GW. Midbrain dopaminergic neurons: a review of the molecular circuitry that regulates their development. Dev Biol. 2013;379:123–38.

88. Lin L, Rao Y, Isacson O. Netrin-1 and slit-2 regulate and direct neurite growth of ventral midbrain dopaminergic neurons. Mol Cell Neurosci. 2005;28:547–55.

89. Karney-Grobe S, Russo A, Frey E, Milbrandt J, DiAntonio A. HSP90 is a chaperone for DLK and is required for axon injury signaling. Proc Natl Acad Sci U S A. 2018;115:E9899–908.

90. Zuehlke AD, Beebe K, Neckers L, Prince T. Regulation and function of the human HSP90AA1 gene. Gene. 2015;570:8–16.

91. Rousseaux MW, de Haro M, Lasagna-Reeves CA, De Maio A, Park J, Jafar-Nejad P, et al. TRIM28 regulates the nuclear accumulation and toxicity of both alpha-synuclein and tau. Elife [Internet]. 2016;5. Available from: 10.7554/eLife.19809

92. Rousseaux MW, Revelli J-P, Vázquez-Vélez GE, Kim J-Y, Craigen E, Gonzales K, et al. Depleting Trim28 in adult mice is well tolerated and reduces levels of α-synuclein and tau. Elife [Internet]. 2018;7. Available from: 10.7554/eLife.36768

93. Villaescusa JC, Li B, Toledo EM, Rivetti di Val Cervo P, Yang S, Stott SR, et al. A PBX1 transcriptional network controls dopaminergic neuron development and is impaired in Parkinson’s disease. EMBO J. 2016;35:1963–78.

94. Pereira Luppi M, Azcorra M, Caronia-Brown G, Poulin J-F, Gaertner Z, Gatica S, et al. Sox6 expression distinguishes dorsally and ventrally biased dopamine neurons in the substantia nigra with distinctive properties and embryonic origins. Cell Rep. 2021;37:109975.

95. Bazargani N, Attwell D. Astrocyte calcium signaling: the third wave. Nat Neurosci. 2016;19:182–9.

96. Zhou Z, Okamoto K, Onodera J, Hiragi T, Andoh M, Ikawa M, et al. Astrocytic cAMP modulates memory via synaptic plasticity. Proc Natl Acad Sci U S A [Internet]. 2021;118. Available from: 10.1073/pnas.2016584118

97. Vardjan N, Kreft M, Zorec R. Dynamics of β-adrenergic/cAMP signaling and morphological changes in cultured astrocytes. Glia. 2014;62:566–79.

98. Henneberger C, Papouin T, Oliet SHR, Rusakov DA. Long-term potentiation depends on release of D-serine from astrocytes. Nature. 2010;463:232–6.

99. Bonneh-Barkay D, Wang G, Starkey A, Hamilton RL, Wiley CA. In vivo CHI3L1 (YKL-40) expression in astrocytes in acute and chronic neurological diseases. J Neuroinflammation. 2010;7:34.

100. Matute-Blanch C, Brito V, Midaglia L, Villar LM, Garcia-Diaz Barriga G, Guzman de la Fuente A, et al. Inflammation in multiple sclerosis induces a specific reactive astrocyte state driving non-cell-autonomous neuronal damage. Clin Transl Med. 2022;12:e837.

101. Buser JR, Maire J, Riddle A, Gong X, Nguyen T, Nelson K, et al. Arrested preoligodendrocyte maturation contributes to myelination failure in premature infants. Ann Neurol. 2012;71:93–109.

102. Akiyama H, Tooyama I, Kawamata T, Ikeda K, McGeer PL. Morphological diversities of CD44 positive astrocytes in the cerebral cortex of normal subjects and patients with Alzheimer’s disease. Brain Res. 1993;632:249–59.

103. Zhu X, Rottkamp CA, Boux H, Takeda A, Perry G, Smith MA. Activation of p38 kinase links tau phosphorylation, oxidative stress, and cell cycle-related events in Alzheimer disease. J Neuropathol Exp Neurol. 2000;59:880–8.

104. Reynolds CH, Nebreda AR, Gibb GM, Utton MA, Anderton BH. Reactivating kinase/p38 phosphorylates tau protein in vitro. J Neurochem. 1997;69:191–8.

105. Zarubin T, Han J. Activation and signaling of the p38 MAP kinase pathway. Cell Res. 2005;15:11–8.

106. Chen J, Ren Y, Gui C, Zhao M, Wu X, Mao K, et al. Phosphorylation of Parkin at serine 131 by p38 MAPK promotes mitochondrial dysfunction and neuronal death in mutant A53T α-synuclein model of Parkinson’s disease. Cell Death Dis. 2018;9:700.

107. Kang W, Balordi F, Su N, Chen L, Fishell G, Hébert JM. Astrocyte activation is suppressed in both normal and injured brain by FGF signaling. Proc Natl Acad Sci U S A. 2014;111:E2987–95.

108. Pringle NP, Yu W-P, Howell M, Colvin JS, Ornitz DM, Richardson WD. Fgfr3 expression by astrocytes and their precursors: evidence that astrocytes and oligodendrocytes originate in distinct neuroepithelial domains. Development. 2003;130:93–102.

109. Fearnley JM, Lees AJ. Ageing and Parkinson’s disease: substantia nigra regional selectivity. Brain. 1991;114 (Pt 5):2283–301.

110. Ris MM, Deitrich RA, Von Wartburg JP. Inhibition of aldehyde reductase isoenzymes in human and rat brain. Biochem Pharmacol. 1975;24:1865–9.

111. Liu Y, Deng S, Song Z, Zhang Q, Guo Y, Yu Y, et al. MLIF Modulates Microglia Polarization in Ischemic Stroke by Targeting eEF1A1. Front Pharmacol. 2021;12:725268.

112. Haure-Mirande J-V, Audrain M, Ehrlich ME, Gandy S. Microglial TYROBP/DAP12 in Alzheimer’s disease: Transduction of physiological and pathological signals across TREM2. Mol Neurodegener. 2022;17:55.

113. Wang N, Wang M, Jeevaratnam S, Rosenberg C, Ikezu TC, Shue F, et al. Opposing effects of apoE2 and apoE4 on microglial activation and lipid metabolism in response to demyelination. Mol Neurodegener. 2022;17:75.

114. Hou J, Chen Y, Grajales-Reyes G, Colonna M. TREM2 dependent and independent functions of microglia in Alzheimer’s disease. Mol Neurodegener. 2022;17:84.

115. Rachmian N, Medina S, Cherqui U, Akiva H, Deitch D, Edilbi D, et al. Identification of senescent, TREM2-expressing microglia in aging and Alzheimer’s disease model mouse brain. Nat Neurosci [Internet]. 2024; Available from: 10.1038/s41593-024-01620-8

116. Keren-Shaul H, Spinrad A, Weiner A, Matcovitch-Natan O, Dvir-Szternfeld R, Ulland TK, et al. A Unique Microglia Type Associated with Restricting Development of Alzheimer’s Disease. Cell. 2017;169:1276–90.e17.

117. Namekata K, Guo X, Kimura A, Arai N, Harada C, Harada T. DOCK8 is expressed in microglia, and it regulates microglial activity during neurodegeneration in murine disease models. J Biol Chem. 2019;294:13421–33.

118. Liu L, Cui Y, Chang Y-Z, Yu P. Ferroptosis-related factors in the substantia nigra are associated with Parkinson’s disease. Sci Rep. 2023;13:15365.

119. Styrpejko DJ, Cuajungco MP. Transmembrane 163 (TMEM163) Protein: A New Member of the Zinc Efflux Transporter Family. Biomedicines [Internet]. 2021;9. Available from: 10.3390/biomedicines9020220

120. Zhao Y, Zhang K, Pan H, Wang Y, Zhou X, Xiang Y, et al. Genetic Analysis of Six Transmembrane Protein Family Genes in Parkinson’s Disease in a Large Chinese Cohort. Front Aging Neurosci. 2022;14:889057.

121. Vela D. The Dual Role of Hepcidin in Brain Iron Load and Inflammation. Front Neurosci. 2018;12:740.

122. Wu K-C, Liou H-H, Kao Y-H, Lee C-Y, Lin C-J. The critical role of Nramp1 in degrading α-synuclein oligomers in microglia under iron overload condition. Neurobiol Dis. 2017;104:61–72.

123. Takeuchi K, Yoshioka N, Higa Onaga S, Watanabe Y, Miyata S, Wada Y, et al. Chondroitin sulphate N-acetylgalactosaminyl-transferase-1 inhibits recovery from neural injury. Nat Commun. 2013;4:2740.

124. Cevik M, Gunduz MK, Deliorman G, Susleyici B. Alterations in niban gene expression as a response to stress conditions in 3T3-L1 adipocytes. Mol Biol Rep. 2020;47:9399–408.

125. Guo X, Li T, Xu Y, Xu X, Zhu Z, Zhang Y, et al. Increased levels of Gab1 and Gab2 adaptor proteins skew interleukin-4 (IL-4) signaling toward M2 macrophage-driven pulmonary fibrosis in mice. J Biol Chem. 2017;292:14003–15.

126. Zheng Y, An H, Yao M, Hou J, Yu Y, Feng G, et al. Scaffolding adaptor protein Gab1 is required for TLR3/4- and RIG-I-mediated production of proinflammatory cytokines and type I IFN in macrophages. J Immunol. 2010;184:6447–56.

127. Tentillier N, Etzerodt A, Olesen MN, Rizalar FS, Jacobsen J, Bender D, et al. Anti-Inflammatory Modulation of Microglia via CD163-Targeted Glucocorticoids Protects Dopaminergic Neurons in the 6-OHDA Parkinson’s Disease Model. J Neurosci. 2016;36:9375–90.

128. Ahuja M, Ammal Kaidery N, Attucks OC, McDade E, Hushpulian DM, Gaisin A, et al. Bach1 derepression is neuroprotective in a mouse model of Parkinson’s disease. Proc Natl Acad Sci U S A [Internet]. 2021;118. Available from: 10.1073/pnas.2111643118

129. Pradhan P, Vijayan V, Cirksena K, Buettner FFR, Igarashi K, Motterlini R, et al. Genetic BACH1 deficiency alters mitochondrial function and increases NLRP3 inflammasome activation in mouse macrophages. Redox Biol. 2022;51:102265.

130. Chen C-M, Wu Y-R, Hu F-J, Chen Y-C, Chuang T-J, Cheng Y-F, et al. HSPA5 promoter polymorphisms and risk of Parkinson’s disease in Taiwan. Neurosci Lett. 2008;435:219–22.

131. Enogieru AB, Omoruyi SI, Hiss DC, Ekpo OE. GRP78/BIP/HSPA5 as a Therapeutic Target in Models of Parkinson’s Disease: A Mini Review. Adv Pharmacol Sci. 2019;2019:2706783.

132. Traag VA, Waltman L, van Eck NJ. From Louvain to Leiden: guaranteeing well-connected communities. Sci Rep. 2019;9:5233.

133. Wang M, Ye R, Barron E, Baumeister P, Mao C, Luo S, et al. Essential role of the unfolded protein response regulator GRP78/BiP in protection from neuronal apoptosis. Cell Death Differ. 2010;17:488–98.

134. Wang M, Wey S, Zhang Y, Ye R, Lee AS. Role of the unfolded protein response regulator GRP78/BiP in development, cancer, and neurological disorders. Antioxid Redox Signal. 2009;11:2307– 16.

135. Dai DL, Li M, Lee EB. Human Alzheimer’s disease reactive astrocytes exhibit a loss of homeostastic gene expression. Acta Neuropathol Commun. 2023;11:127.

136. Sosunov A, Wu X, McGovern R, Mikell C, McKhann GM 2nd, Goldman JE. Abnormal mitosis in reactive astrocytes. Acta Neuropathol Commun. 2020;8:47.

137. van Abel D, Abdulhamid O, Scheper W, van Dijk M, Oudejans CBM. STOX1A induces phosphorylation of tau proteins at epitopes hyperphosphorylated in Alzheimer’s disease. Neurosci Lett. 2012;528:104–9.

138. Partanen J, Achim K. Neurons gating behavior-developmental, molecular and functional features of neurons in the Substantia Nigra pars reticulata. Front Neurosci. 2022;16:976209.

139. Liu D, Li W, Ma C, Zheng W, Yao Y, Tso CF, et al. A common hub for sleep and motor control in the substantia nigra. Science. 2020;367:440–5.

140. Terkelsen MH, Hvingelby VS, Pavese N. Molecular Imaging of the GABAergic System in Parkinson’s Disease and Atypical Parkinsonisms. Curr Neurol Neurosci Rep. 2022;22:867–79.

141. Alberico SL, Cassell MD, Narayanan NS. The Vulnerable Ventral Tegmental Area in Parkinson’s Disease. Basal Ganglia. 2015;5:51–5.

142. Graybiel AM, Ohta K, Roffler-Tarlov S. Patterns of cell and fiber vulnerability in the mesostriatal system of the mutant mouse weaver. I. Gradients and compartments. J Neurosci. 1990;10:720–33.

143. Triarhou LC, Norton J, Ghetti B. Mesencephalic dopamine cell deficit involves areas A8, A9 and A10 in weaver mutant mice. Exp Brain Res. 1988;70:256–65.

144. Philippens IHCHM, Wubben JA, Franke SK, Hofman S, Langermans JAM. Involvement of the Red Nucleus in the Compensation of Parkinsonism may Explain why Primates can develop Stable Parkinson’s Disease. Sci Rep. 2019;9:880.

145. Halliday G, Reyes S, Double K. Chapter 13 - Substantia Nigra, Ventral Tegmental Area, and Retrorubral Fields. In: Mai JK, Paxinos G, editors. The Human Nervous System (Third Edition). San Diego: Academic Press; 2012. p. 439–55.

146. Beine Z, Wang Z, Tsoulfas P, Blackmore MG. Single Nuclei Analyses Reveal Transcriptional Profiles and Marker Genes for Diverse Supraspinal Populations. J Neurosci. 2022;42:8780–94.

147. Hough HB, Wolff HG. The relative vascularity of subcortical ganglia of the cat’s brain; the putamen, globus pallidus, substantia nigra, red nucleus, and geniculate bodies. J Comp Neurol. 1939;71:427–36.

148. Huisman M. Collateralization of descending spinal pathways from red nucleus and other brainstem cell groups in rat, cat and monkey. 1983; Available from: https://repub.eur.nl/pub/51247/

149. Jiang J, Wang C, Qi R, Fu H, Ma Q. scREAD: A Single-Cell RNA-Seq Database for Alzheimer’s Disease. iScience. 2020;23:101769.

150. Qin Y, Qiu J, Wang P, Liu J, Zhao Y, Jiang F, et al. Impaired autophagy in microglia aggravates dopaminergic neurodegeneration by regulating NLRP3 inflammasome activation in experimental models of Parkinson’s disease. Brain Behav Immun. 2021;91:324–38.

151. Lv Q-K, Tao K-X, Wang X-B, Yao X-Y, Pang M-Z, Liu J-Y, et al. Role of α-synuclein in microglia: autophagy and phagocytosis balance neuroinflammation in Parkinson’s disease. Inflamm Res. 2023;72:443–62.

152. Tan JX, Finkel T. Lysosomes in senescence and aging. EMBO Rep. 2023;24:e57265.

153. Nixon RA. The aging lysosome: An essential catalyst for late-onset neurodegenerative diseases. Biochim Biophys Acta: Proteins Proteomics. 2020;1868:140443.

154. Norden DM, Godbout JP. Review: microglia of the aged brain: primed to be activated and resistant to regulation. Neuropathol Appl Neurobiol. 2013;39:19–34.

155. Perry VH, Holmes C. Microglial priming in neurodegenerative disease. Nat Rev Neurol. 2014;10:217–24.

156. Guo S, Wang H, Yin Y. Microglia Polarization From M1 to M2 in Neurodegenerative Diseases. Front Aging Neurosci. 2022;14:815347.

157. Jurga AM, Paleczna M, Kuter KZ. Overview of General and Discriminating Markers of Differential Microglia Phenotypes. Front Cell Neurosci. 2020;14:198.

158. Javanmehr N, Saleki K, Alijanizadeh P, Rezaei N. Microglia dynamics in aging-related neurobehavioral and neuroinflammatory diseases. J Neuroinflammation. 2022;19:273.

159. Conner SC, Benayoun L, Himali JJ, Adams SL, Yang Q, DeCarli C, et al. Methionine Sulfoxide Reductase-B3 Risk Allele Implicated in Alzheimer’s Disease Associates with Increased Odds for Brain Infarcts. J Alzheimers Dis. 2019;68:357–65.

160. Beach TG, Sue LI, Walker DG, Lue LF, Connor DJ, Caviness JN, et al. Marked microglial reaction in normal aging human substantia nigra: correlation with extraneuronal neuromelanin pigment deposits. Acta Neuropathol. 2007;114:419–24.

